# Positive selection on hotspot and reinforcing regulatory alleles contributed to hexaploid bread wheat improvement

**DOI:** 10.1101/2025.10.08.680093

**Authors:** Shuhua Zhan, Elie Raherison, William Hargreaves, Nia Hughes, Roos Goessen, Mohammad Mahdi Majidi, Ron Knox, Richard Cuthbert, Lewis Lukens

## Abstract

**Background:** Genetic variation of regulatory alleles plays a key role in evolution and breeding. In polyploids, regulatory differences may preferentially affect genes on homoeologous chromosomes or sub-genomes. Selection in plant breeding may act upon total transcript dosage across homoeologous genes and on alleles that have strong effects on the transcriptome.

**Results:** To investigate these questions, we identified regulatory polymorphisms between an old and a recent hexaploid bread wheat cultivar (*Triticum aestivum*, 2n=6x=42, AABBDD). The recent cultivar was the product of decades of selection for grain yield and quality. Regulatory allele polymorphisms preferentially affected genes on homoeologous chromosomes but rarely affected genes on specific sub-genomes. The chromosomal distributions of regulatory alleles indicated that past selection had acted upon them, and the effect of selection differed between alleles targeting environmental response genes and genes involved in other processes. Modern cultivar alleles that affected many genes’ transcripts corresponded to known selection targets and improved field crop performance. Modern cultivar alleles also had significant effects on homoeologous genes, and these alleles also improved crop performance.

**Conclusions:** Polyploid breeding across many species has been and will continue to be the key factor in plant improvement. By enhancing the favorability of strong regulatory alleles and by expanding the range of gene transcript abundances, genome duplications enable breeding progress.

## Background

Increasing the supply of cereal grain as well as other plant products is critical to meet the demands of a growing population and to reduce the environmental impact of agriculture. Modern plant breeding works to improve current high-yielding genotypes adapted for specific growing environments. Plant breeding has quickly improved traits in a wide range of species [1, 2]. For example, Canadian and European wheat cultivar grain yields have improved between 38 kg ha^-1^ year^-1^ to 128 kg ha^-1^ year^-1^ in the past century [1, 3, 4].

Regulatory loci that affect gene transcript abundances have contributed to desired traits in plant improvement. Domestication selects for the traits necessary for agricultural production [5] and often involves large effect regulatory alleles [6]. Adaptation selects for the traits necessary to cultivate in a region such as a short maturity time and tolerance to specific abiotic and biotic stresses. Breeding typically selects among the progeny of adapted line crosses after rigorous evaluation [5]. Adaptation and breeding mostly occur through the contributions of many small effect alleles, although major regulatory alleles play a role [7, 8]. Rapid evolution in natural populations occurs through regulatory changes in ways analogous to crop adaptation and breeding processes [9–11].

Many crop species are polyploids [12]. Polyploid regulatory alleles function in the context of sub-genomes. Allelic differences at trans-acting regulatory loci may affect genes on homoeologous chromosomes. Although inter-homeolog regulation can be complex [13], Bao et al. 2019, He et al. 2022 and Tan et al. 2022 reported cases of putative genetically induced feedback regulation in which a regulatory allele that upregulated a gene downregulated its homeolog(s) [14–16]. In addition, regulatory polymorphisms may cause sub-genome specific transcriptional responses. Genes on specific sub-genomes have been reported to be co-regulated. In tetraploid *Coffea arabica* exposed to water deficit, salt stress, and high temperature, the transcriptional response of mannitol biosynthesis and catabolism genes differed across sub-genomes [17]. More genes on the bread wheat B and D sub-genomes responded to pathogen attack than did genes on the A sub-genome, and the transcriptional changes of the B and D genes were higher than the A genes [18].

Selection on regulatory alleles may differ between polyploids and diploids. While a novel regulatory allele frequently affects a fitness-related trait in a diploid, a novel allele in a polyploid may have little or no effect on traits because homeologs mask its effects. Mutagenesis is much more likely to generate a noticeable phenotype in the diploid grass barley than in its hexaploid wheat relative [19]. However, homeolog masking may not be complete. Hexaploid wheat’s homoeologous genes have low allelic variation at nonsynonymous sites [20, 21] suggesting that mutations that affected single gene functions also affected plant fitness.

Selection may more likely favor derived allelic variants that affect numerous genes, hotspot alleles, in cultivated polyploid populations than in diploid populations. Hotspot alleles are broadly deleterious [11, 22, 23], but mutations that natural selection would eliminate in ancestral environments may be neutral or positive in plant breeding [21]. In addition, homeologs may maintain the ancestral functions lost by hotspot alleles [19, 24]. Selection has acted upon several, large-effect dominant and semi-dominant genes that may be hotspots during wheat adaptation and advancement [25].

Homoeologous loci can also together favorably contribute to traits [26, 27]. Accumulating alleles that consistently regulate homoeologous genes can enhance or reduce traits affected by metabolic pathway flux [28]. In bread wheat, delayed leaf senescence may be favorable. Loss of function mutants in the A and D sub-genome homeologs of the Gpc-B1 gene individually led to delayed senescence. Senescence was even more delayed in the double mutant [27].

In this work, we identified and analyzed regulatory allele polymorphisms between an older bread wheat cultivar, Red Fife, dominant in Canada in the early 1900s and a recently developed variety Stettler that has notably higher yields and yield stability [29]. The first objective of this work was to investigate how hexaploid bread wheat regulatory locus polymorphisms affected homoeologous genes and genes on specific sub-genomes. The second objective was to investigate if selection frequently affected regulatory alleles. The third and final objective was to determine if plant breeders selected hotspot alleles and reinforcing regulatory alleles.

## Results

### Characterizing regulatory polymorphisms and their target genes within polyploid wheat

#### Many regulatory alleles differ between closely related Red Fife and Stettler cultivars

To identify the regulatory allele differences between the ancestral and modern cultivars, we measured gene transcript abundances from a population of 154 homozygous doubled haploid lines (DHLs) developed from Red Fife x Stettler F1 microspores. From the genome’s 110,790 total annotated high-confidence genes, 40,395 (36.5%) were expressed in leaf tissue, and the 30,296 (75.0%) most variable expressed genes were selected to identify eQTL. Expressed genes were located on the A, B, and D sub-genomes at similar frequencies **(Fig. 1)**. Each sub-genome had a similar proportion of variable genes **(Fig. 1)**.

**Fig. 1.**
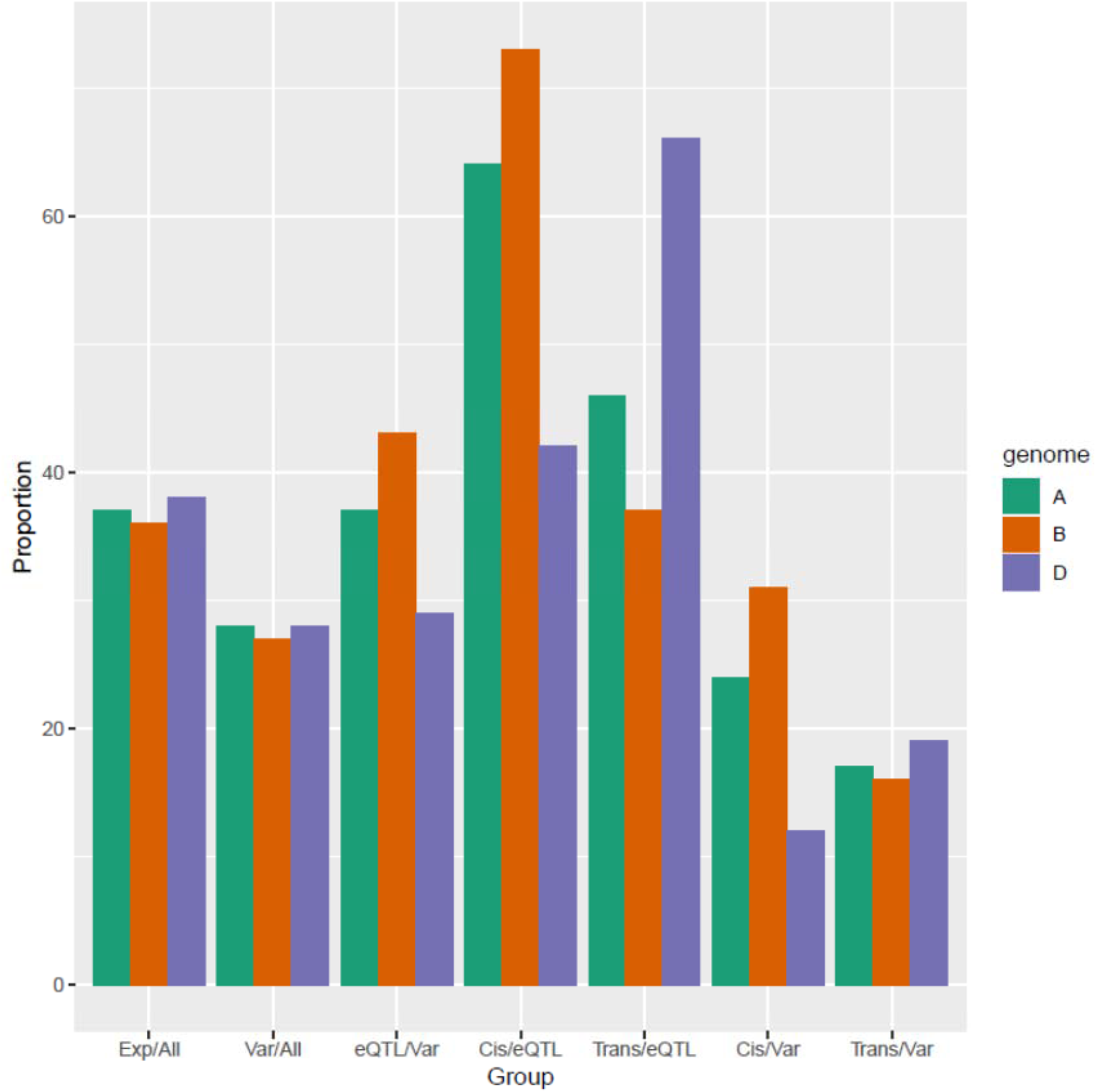
Proportions of expressed, variable, and eQTL regulated genes amongst different gene sets across the A, B, and D subgenomes. Exp/All and Var/All are, respectively, the proportions of annotated genes on sub-genomes that are expressed and are within the top 75.0% variable genes. eQTL/Var is the proportion of variable genes regulated by eQTL. Cis/eQTL and Trans/eQTL are the proportions of eQTL regulated genes that are cis- and trans-eQTL regulated, respectively. Cis/Var and Trans/Var are the proportions of all variable genes that have cis-eQTL and trans-eQTL.

A total of 12,514 eQTL significantly contributed to the expression variation of 10,949 genes, 36.1% of the variable genes (**Additional file 1: Table S1**). While this study’s population type and experimental design provided a high power to detect eQTL, small effect regulatory loci below detection limits also very likely contributed to expression differences. Gene expression variation was partitioned into genetic sources and unexplained environmental sources. We estimated the environmental variance for each expressed gene without a detected eQTL using the two sets of four replicated, homozygous parental lines. Genes’ expression variances across the DHL population were consistently greater than their estimated environmental variances. Among the 29,446 genes for which we did not detect eQTL, 25,345 (86.1%) had lower variances across the replicated genotypes than across different genotypes.

#### Polymorphic trans acting regulatory loci usually affected several genes and had smaller effects on a gene’s transcript abundance than did cis regulatory loci

We classified eQTL as cis-acting loci, corresponding to regulatory variants linked to or within their associated genes, and trans acting loci, corresponding to regulatory variants unlinked to their associated genes. A gene’s cis-eQTL variant may be within a promoter or an enhancer element that affects the gene. A gene’s trans-eQTL variant may be within the regulatory or coding sequence of a regulatory gene or of an enzyme-encoding gene with downstream effects on other genes. The cis regulatory loci slightly outnumbered trans regulatory loci. Of all eQTL, 6,636 (53.0%) were cis-eQTL and 5,878 (47.0%) were trans-eQTL (**Additional file 1: Table S1**). More genes were cis-regulated (60.6%; 6,634/10,949 genes because two genes were each regulated by two cis-eQTL) than trans regulated (48.7%; 5,328/10,949 genes) (**Additional file 1: Table S2**). The sum of the two percentages was more than 100 because both cis- and trans-eQTL regulated 1,013 (9.3%) of the 10,949 genes. A total of 5,621 genes (51.3%) were regulated by only cis-eQTL, and 4,315 genes (39.4%) were regulated by only trans-eQTL (**Additional file 1: Table S2, Fig. 1**).

Both cis- and trans-eQTL had strong effects on gene expression, and cis-eQTL effects were larger than trans-eQTL effects. The proportions of transcript abundance variation explained by each of the 6,636 cis-eQTL were close to uniformly distributed with an average of 46.4% (**Fig. 2A**). Each of the 5,878 trans-eQTL explained, on average, 24.0% of a target gene’s expression variation. The trans-eQTL R^2^ values were strongly skewed to lower proportions, and a high proportion of the trans-eQTL explained less than 24% of gene expression variance (**Fig. 2A**).

**Fig. 2.**
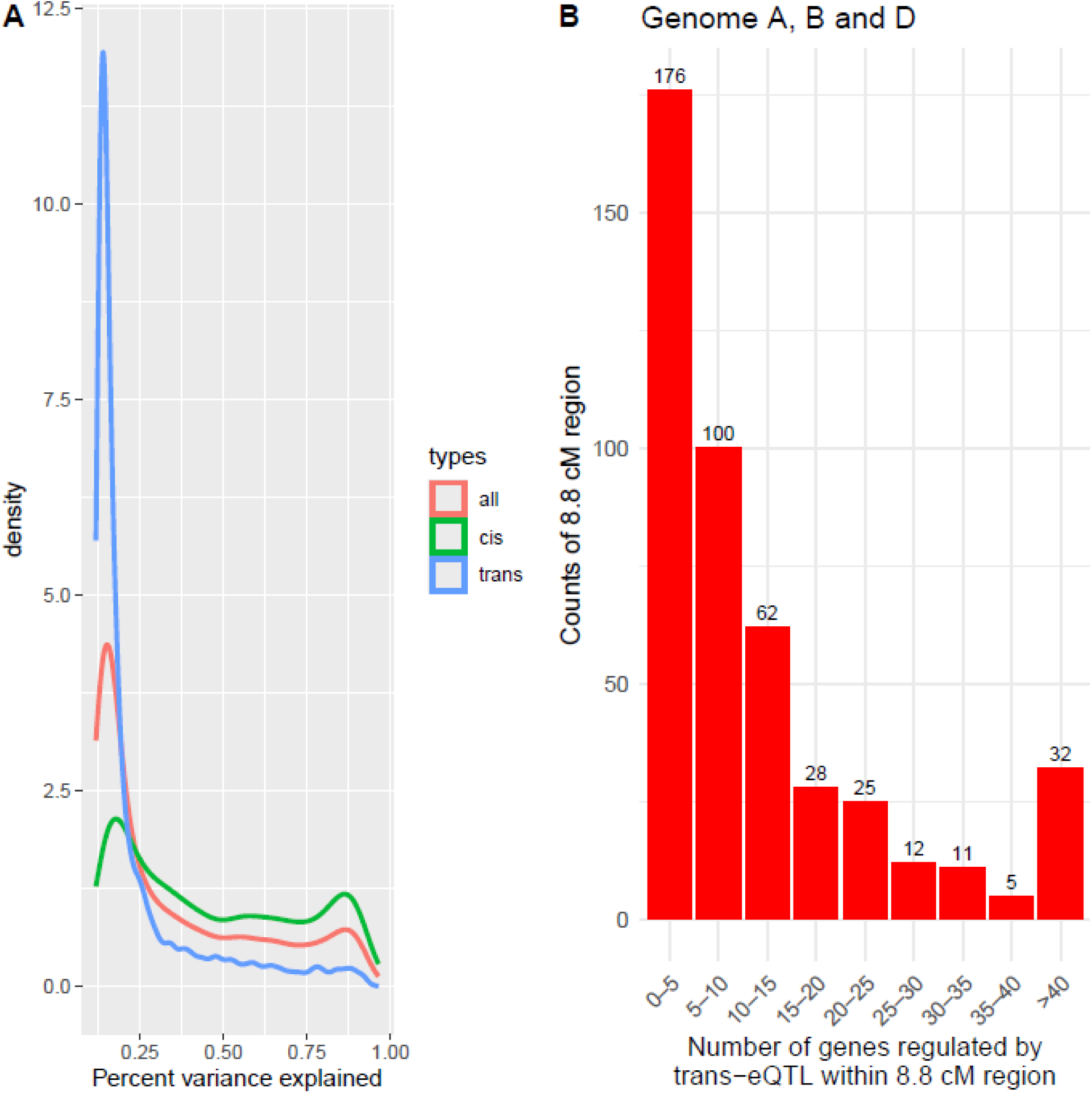
**A** A density plot of the percent of a gene’s expression variance explained by an eQTL, R^2^. The figure shows the R^2^ distributions for all QTL, cis-eQTL, and trans-eQTL. **B** The number of regulatory intervals that regulate from zero to more than 40 genes.

Most trans-acting chromosomal regions regulated a few genes although several regions regulated many, and different trans-eQTL often affected the same genes. Across the genome, 42 loci were regulatory hotspots, short genetic intervals dense with eQTL (**Additional file 1: Table S3**). Of 451 8.8cM genetic map intervals, 399 contained one or more trans eQTL. The number of genes regulated by an interval had first quartile, median, and third quartile values of 3.5, 8, and 17, respectively (**Fig. 2B**). Some genes regulated by a single interval were likely regulated by different loci. Trans regulatory alleles, as well as cis regulatory alleles, were notably less frequent in the D genome, and the number of genes regulated by the 104 D genome intervals with at least one eQTL had first quartile, median, and third quartile values of 2, 5, and 12 (**Additional file 2: Fig. S1**). Different trans-eQTL intervals regulated overlapping sets of genes. While most trans-eQTL regulated several genes, the number of trans-eQTL gene targets, 5,328, was less than the number of trans-eQTL, 5,878.

#### Almost all trans-eQTL gene targets were uniformly distributed across sub-genomes, and many targets were preferentially located on homoeologous chromosomes

With few exceptions, a trans-eQTL’s gene targets were located at equal frequencies in the three sub-genomes. Excluding gene targets on homoeologous chromosomes, trans-eQTL targeted genes were in different sub-genomes at close to twice the frequency as they were on the same sub-genome, (64.9% vs. 35.1%; X^2^ test, p-value = 0.143; **Fig. 3A; Additional file 1: Table S4**). We also examined groups of co-regulated genes for sub-genome enrichment. A total of 119 regulatory intervals affected 10 or more genes on non-homoeologous chromosomes. Only four of these intervals, 3.36%, significantly affected genes located one sub-genome (FDR, false discovery rate, p value <0.05) (**Additional file 1: Table S5**). These four intervals targeted genes were enriched for specific biological processes (**Additional file 1: Table S6**).

**Fig. 3.**
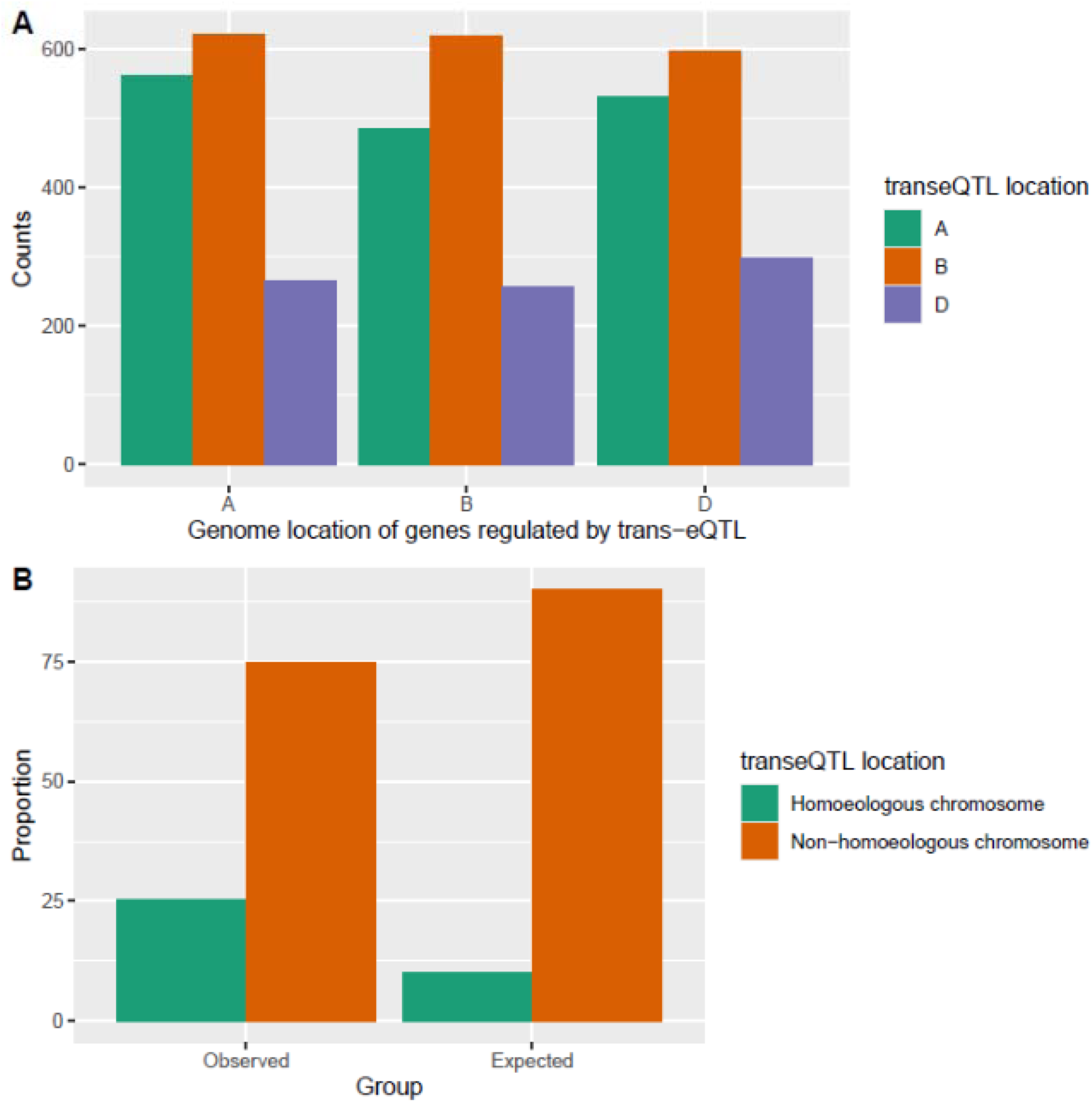
The locations of trans eQTL target genes. **A** The numbers of genes on the A, B, and D genomes regulated by trans-eQTL on the A, B and D genomes are shown. **B** The expected and observed proportions of genes’ trans-eQTL that were on homoeologous or non-homoeologous chromosomes.

Trans-eQTL on one chromosome preferentially regulated genes located on homoeologous chromosomes. Twenty-five percent of the genes regulated by trans-eQTL were regulated by a locus on a homoeologous chromosome, notably higher than the expected ∼10% (**Fig. 3B; Additional file 1: Table S7**; one-sample proportions test, p-value < 0.001). The proportion of genes regulated by homoeologous trans-eQTL differed among chromosomes. For example, 40.6% of the trans-eQTL that regulated chromosome 2A genes were located on linkage groups 2B and 2D while 8.3% of trans-eQTL that regulated chromosome 7B genes were located on linkage groups 7A and 7D (**Additional file 1: Table S7; Additional file 2: Fig. S2**).

The enrichment of trans-eQTL target genes on homoeologous chromosomes was in part due to cis-eQTL which generated a feedback control type process across homeologs to maintain a uniform level of total homeolog expression. A total of 152 chromosomal intervals each contained both a cis-eQTL for one gene in a triplet and a trans-eQTL for a homeolog in the same triplet. Eight additional intervals contained a cis-eQTL for one gene and two trans-eQTL for the gene’s two homeologs (**Fig. 4**). These co-localized cis and trans alleles usually had opposite effects on their target genes thus contributing to a uniform level of total homeolog expression across homeologs. The 152 intervals differentially regulated the two target genes 4.4 times more often than they consistently regulated them (124 vs 28) (**Fig. 4**). In five of the eight intervals containing a cis-eQTL for one homeolog and a trans-eQTL for each of the two other homeologs, the cis-eQTL and both trans-eQTL had opposite effects (**Fig. 4)**. The same interval accounted for 27.8% of the 605 cases where a cis-eQTL regulated one homeolog and a trans-eQTL regulated another homeolog.

**Fig. 4.**
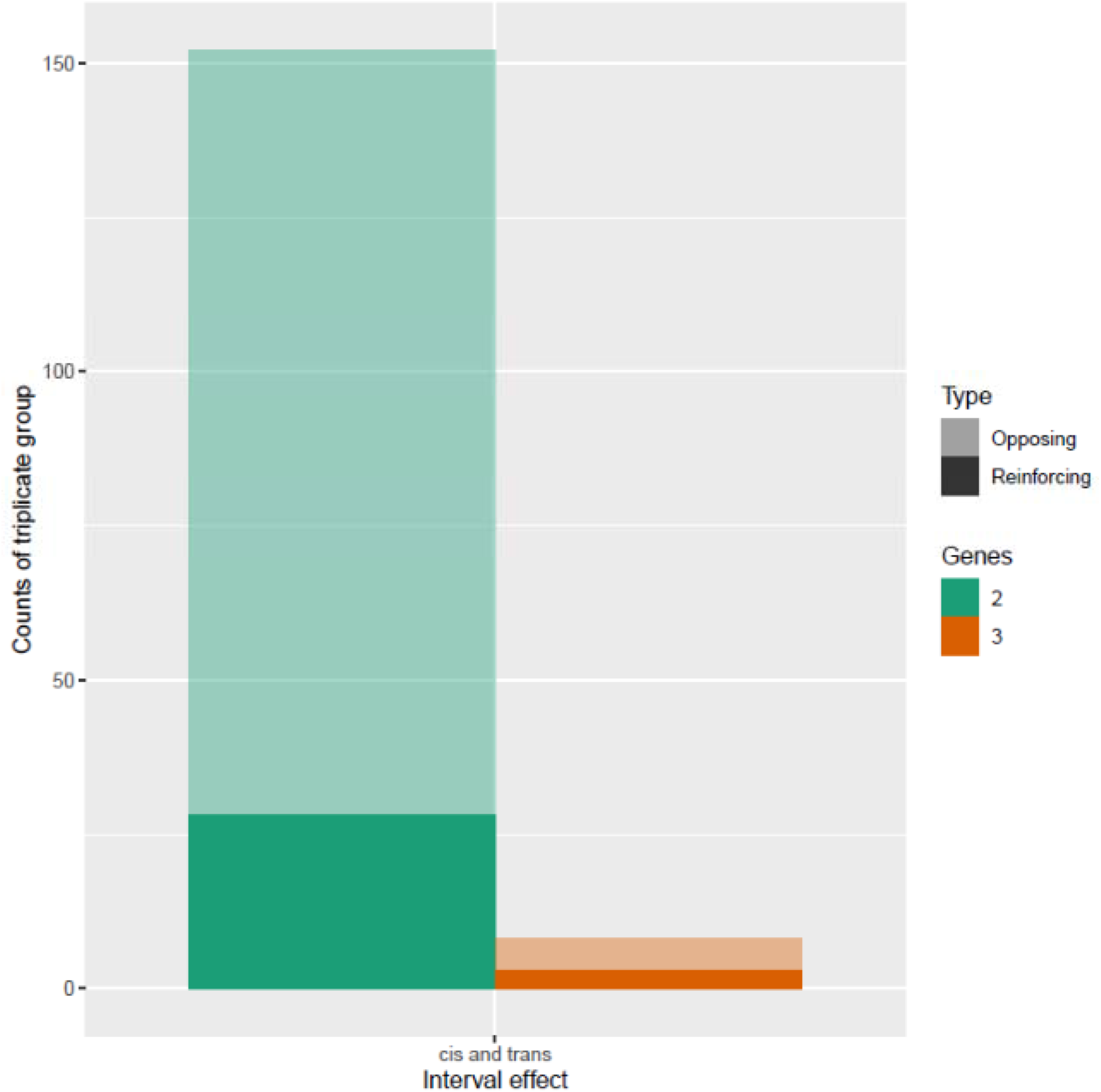
An allele that cis regulates one gene and trans regulates its homeologs often has opposite effects. The number of triplicates in which a chromosomal interval regulated one homeolog in cis and either one or two others in trans are shown. One parent’s alleles consistently regulated homeologs (darker colors) or upregulated one and downregulated the other (lighter colors).

### Selection appears to have acted upon many regulatory alleles and to have affected regulatory alleles differently depending on their target gene functions

We investigated if selection had often acted on regulatory alleles by examining the intra-chromosomal distributions of eQTL. Purifying or positive selection leads to allele fixation or loss, so if eQTL contributed to plant fitness, they would be less frequent in genomic regions where selection had been effective relative to genomic regions where selection had been less effective. Selection is less effective on alleles in chromosome regions with low crossing over [30, 31], and large pericentromeric chromosomal regions had notably suppressed crossing over compared to chromosome ends [21] (**Additional file 2: Fig. S3**). The number of cis-eQTL per synonymous nucleotide difference, an estimate of neutral sequence divergence, in central regions was 2.99, more than twice the number on chromosome ends, 1.48 (paired t test, p-value = 0.045; **Fig. 5**). The number of trans-eQTL per synonymous nucleotide difference was nominally higher (1.58x) in chromosome centers than in chromosome arms (3.15 vs. 1.99 paired t test, p-value = 0.289; **Fig. 5**). Nonsynonymous nucleotide differences were also enriched in central regions, consistent with reduced selection efficacy. The mean ratios of nonsynonymous to synonymous differences were 2.13 for the central regions and 0.98 for the ends (**Fig. 5**, paired t test, p-value = 0.005).

**Fig. 5.**
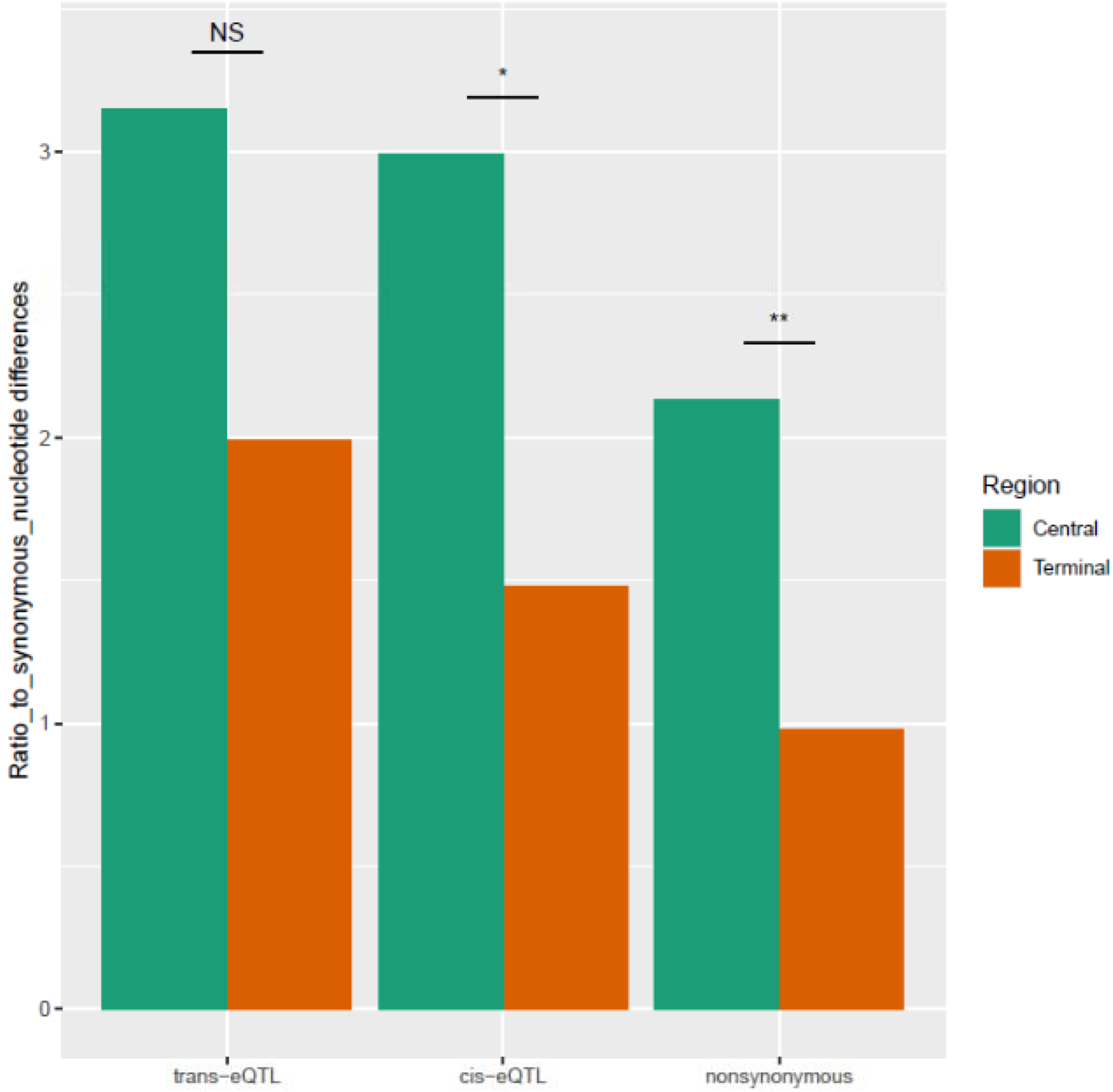
The enrichment of eQTL and nonsynonymous SNPs within low recombination centromeric regions. The x axis gives the feature found in central or terminal chromosome regions. The y axis gives the mean of the number of features corrected for synonymous nucleotide differences. The frequency of features between the terminal regions and central regions significantly differed for cis eQTL and for nonsynonymous SNPs. Symbols: * and ** indicate the corrected feature count across chromosomes was greater in central than in terminal regions at p<0.05, and p<0.01, paired t test, respectively.

If past selection acted similarly on regulatory alleles regardless of their gene targets, genes with different attributes would have a similar probability of being affected by Red Fife and Stettler eQTL. Triplicated genes’ transcript abundances were less variable than non-triplicated genes’, and eQTL were less likely to regulate variable triplicated genes than non-triplicated genes (**Additional file 1: Table S8**, **Additional file 2: Fig. S4**). Trans eQTL affected genes involved in environmentally responsive processes much more often than genes involved in other biological processes. Eighty-one GO biological process (BP) terms were significantly enriched amongst the 5,328 trans-regulated genes (Fisher’s exact test, P<0.01; **Additional file 1: Table S9)**. The top three enriched terms were plant-type hypersensitive response, protein phosphorylation, and response to water deprivation (each with P< 1.0 E-06), and all but two of the most significantly enriched 32 BP GO terms with P values ranging from 2.4 E-07 to 2.75 E-03 were related to environmental response (**Fig. 6A**). Certain genes were also more likely have cis-eQTL. A set of 39 BP terms were enriched in the genes regulated by cis-eQTL at P<0.01 (**Additional file 1: Table S9**). Several enriched terms were environmental response related, but genes related to specific developmental and other processes were also over-represented. **Fig. 6B** shows the 32 most significantly enriched BP terms in the cis-eQTL regulated genes.

**Fig. 6.**
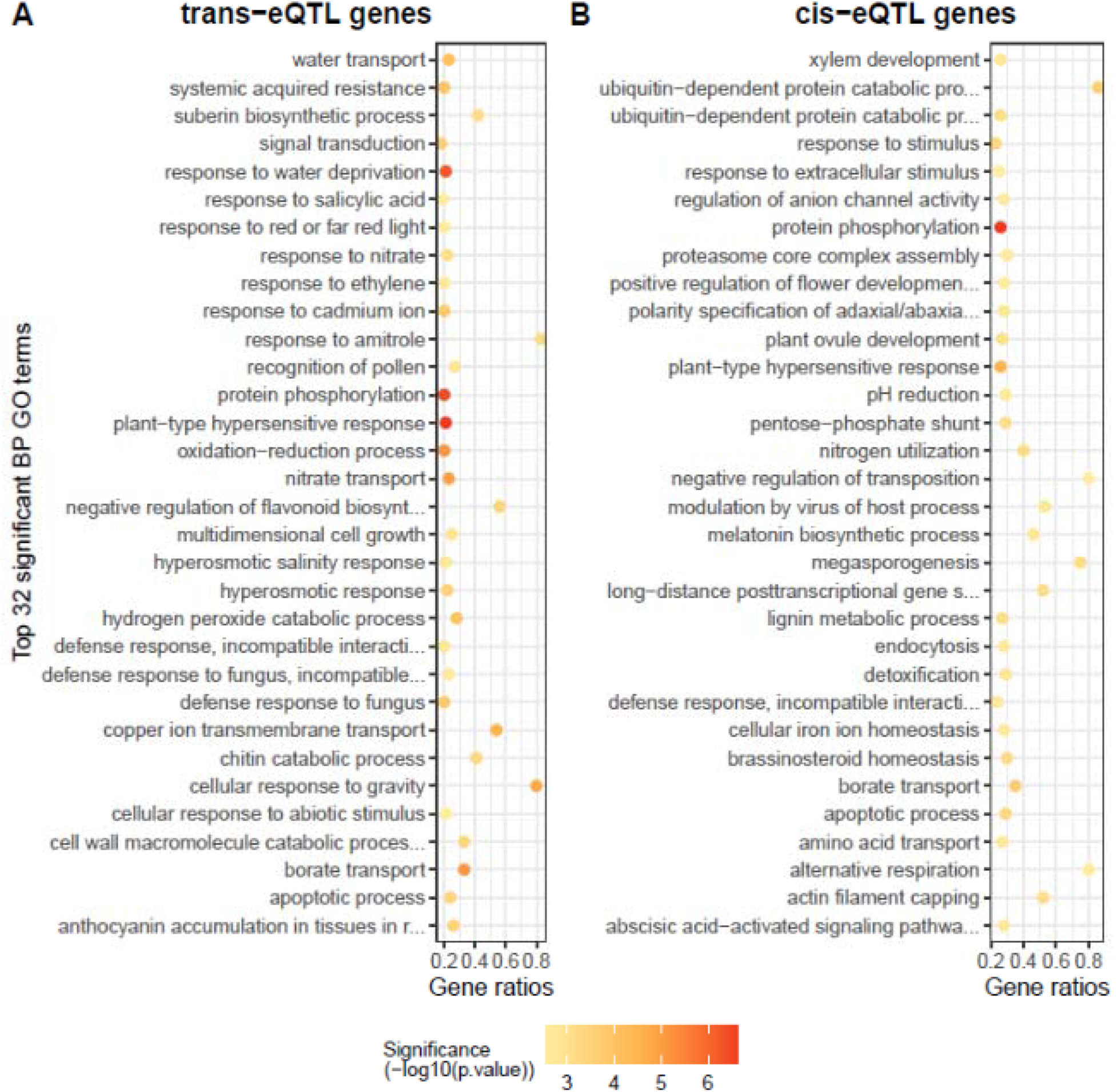
A dotpot of biological process, BP, GO terms enriched amongst genes regulated by cis-eQTL and trans-eQTL. **A** The top 32 enriched BP GO terms out of 81 terms (Fisher’s exact test, P<0.01) enriched amongst genes regulated by trans-eQTL. All terms except multidimensional cell growth and oxidation reduction were considered processes that occur in response to environmental stimuli. **B** The top 32 of 39 enriched (P<0.01) BP GO terms enriched amongst genes regulated by cis-eQTL were involved in more diverse processes. Gene ratios are the proportions of the expressed genes’ GO terms identified within an eQTL group.

### Selection on hotspot alleles and alleles regulating homoeologous genes contributed to improved bread wheat traits

#### Selection for hotspot alleles in the plant breeding era likely contributed to wheat improvement

The regulatory allele enrichment in low recombination chromosome domains indicated past selection. The regulatory alleles that drove cultivar improvement from plant breeding were of specific interest. We determined if genes known to have been selected by plant breeders corresponded to one of the 42 regulatory hotspots. While many major genes in wheat improvement have been identified, two genes, *Lr2a* and *Rht-B1b*, unambiguously differed between Red Fife and Stettler. Both genes corresponded to a regulatory hotspot (**Additional file 1: Table S3**). The *Lr2a* gene [32] confers complete resistance to a large proportion of the stem rust, *Puccinia triticina,* isolates in Canada [33] and corresponded to a hotspot in linkage group 2D, cho2D_83_97. This hotspot regulated 316 genes (**Additional file 1: Table S3**) which were enriched for processes including pathogen response pathways such as salicyclic acid biosynthetic process [34], jasmonic acid mediated signaling pathway [35], and response to ethylene [36] (**Additional file 1: Table S10**). Stettler is homozygous for the semidominant *Rht-B1* “Green Revolution” semi-dwarfing gene. Although a previous report identified no significant transcriptome differences between *Rht-B1b* and wild-type genotypes [37], the locus differentially regulated 34 genes (**Additional file 1: Table S3**). The Stettler *Rht-B1b* hotspot allele upregulated gibberellin 3-beta-hydroxylase 2 transcript. The Stettler *Rht-B1b* allele reduces plants’ responses to GA [38] and increases *in planta* gibberellin levels [39]. Gibberellin 3-beta-dioxygenase, or equivalently gibberellin 3-beta-hydroxylase, converts the inactive gibberellin (GA) precursors GA9 and GA20 into the bioactive gibberellins, GA4 and GA1, respectively [40]. The hotspot also regulated an indole-3-acetic-acid amido synthase gene (TraesCS2A01G277500) that affects cell expansion [41]. Semidominant *Rht-B1b* plants had smaller cell sizes than wild-type plants [42].

Additional hotspot alleles contributed to grain yield, the major breeding target, as well as other favorable traits. The forty-two eQTL hotspot genotypes predicted grain yield, plant height, spike length, and thousand kernel weight (TKW) in field grown plants notably better than did predictions generated by random locus-trait associations (P<0.001, Welch Two Sample t-test; **Additional file 2: Fig. S5**). (Hotspot predictions of heading date, lodging, and protein content were not consistently better than random marker predictions **(Additional file 2: Fig. S5)**). Variable selection identified the specific hotspots that significantly contributed to traits. Genotypes homozygous for different 4B, *Rht-B1* hotspot alleles had a 0.96 standard deviation difference in plant height. Nine hot-spots were strongly associated with TKW, a component of yield (**Fig. 7**). Three contributed to yield, and Stettler alleles at all three had positive effects.

**Fig. 7.**
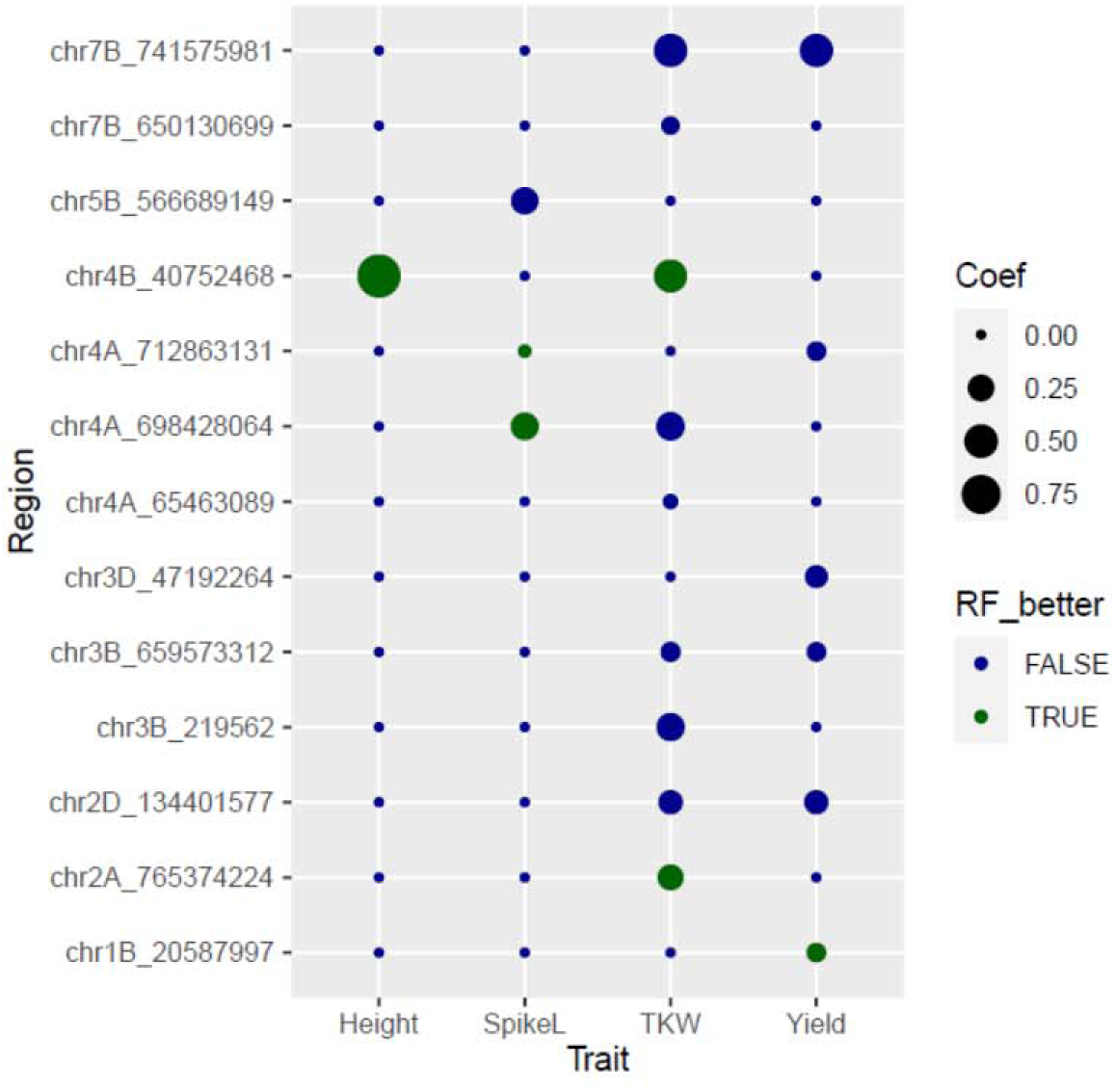
Trans-eQTL hotspots that contributed to trait variation and the magnitudes of their contributions. The hotspots with nonzero coefficients from LASSO regression of trait values onto 42 trans-eQTL hotspots are in the first column. The magnitude of an allelic effect at a hotspot is expressed in standard deviations and represents allelic effects. Coefficients for which the RF allele increases trait values are coloured blue.

Six hotspots contributed to grain-yield. For five of the six hotspots, the Stettler allele had a positive yield contribution (**Fig. 7**). We focused on these hotspot genes’ functions. Genes differentially expressed by distinct hotspots were were biologically coherent and enriched for specific GO biological processes (P<0.01, Fisher’s exact test) including defense response, photosynthesis, energy production, and carbohydrate biosynthesis (**Additional file 1: Table S10**). The most notable yield hotspot was on chr7B. Individuals homozygous for the Stettler allele had yields one standard deviation higher than individuals homozygous for the Red Fife allele and also higher TKW. Biological process terms including energy production and cell division were enriched amongst differentially regulated genes. Of eight genes annotated with “mitochondrial genome maintenance,” two were in this group (P<0.0001; **Additional file 1: Table S10**). A second yield hotspot interval 4A_146_152 regulated genes which were involved in environmental response processes including defense response to nematode, response to endoplasmic reticulum stress, and response to salt stress (P<0.0001). A third hotspot interval 3D_27_29 regulated genes which were enriched for photosynthesis related terms (P<0.0001). A fourth hotspot interval 2D_83_97 regulated genes which were involved starch biosynthetic process, tryptophan catabolic process, isopentenyl diphosphate biosynthetic process, glycogen biosynthetic process, indoleacetic acid biosynthetic process, cellular response to fructose stimulus, and cellular response to sucrose stimulus (P<0.0001).

#### Reinforcing regulatory alleles were over-represented in modern wheat and contributed to wheat yields and other traits

We next evaluated if breeders selected regulatory alleles with the same effects on homeologs. The accumulation of different regulatory alleles that consistently affect gene expression in one lineage is evidence for positive selection on these alleles [8,43], and we tested if Stettler alleles at different regulatory loci had consistent effects on homoeologous genes. Two distinct regulatory loci may affect two homeologs with three types of regulatory patterns. In Type I, two cis-eQTL regulate two homoeologous genes. In Type II, a cis-eQTL regulates one gene, and a trans-eQTL regulates its homeolog. In Type III, two trans-eQTL each regulate a homoeologous gene **(Fig. 8)**. The frequency with which one parent’s cis-eQTL alleles regulated homeologs in the same way was similar to the frequency that they regulated homeologs in opposite ways. Out of a total of 326 Type I cases, in 162 (50.4%) the same parent’s allele at the two regulatory loci consistently affected their target genes (**Fig. 8A**; X^2^ test, p-value = 0.908). Similarly, consistent cis- and trans-eQTL regulation of different homeologs was not significantly enriched amongst Type II cases. Among the 437 Type II cases, 208, 48%, had consistently acting parental alleles. (**Fig. 8B**; X^2^ test, p-value = 0.234). In contrast, for 64% of the Type III cases (93/146), one parent’s alleles at the two trans-eQTL had the same effects on homoeologous genes (**Fig. 8C**; X^2^ test, p-value < 0.001). This enrichment indicated that plant breeders selected separate trans-eQTL to co-regulate homoeologous genes.

**Fig. 8.**
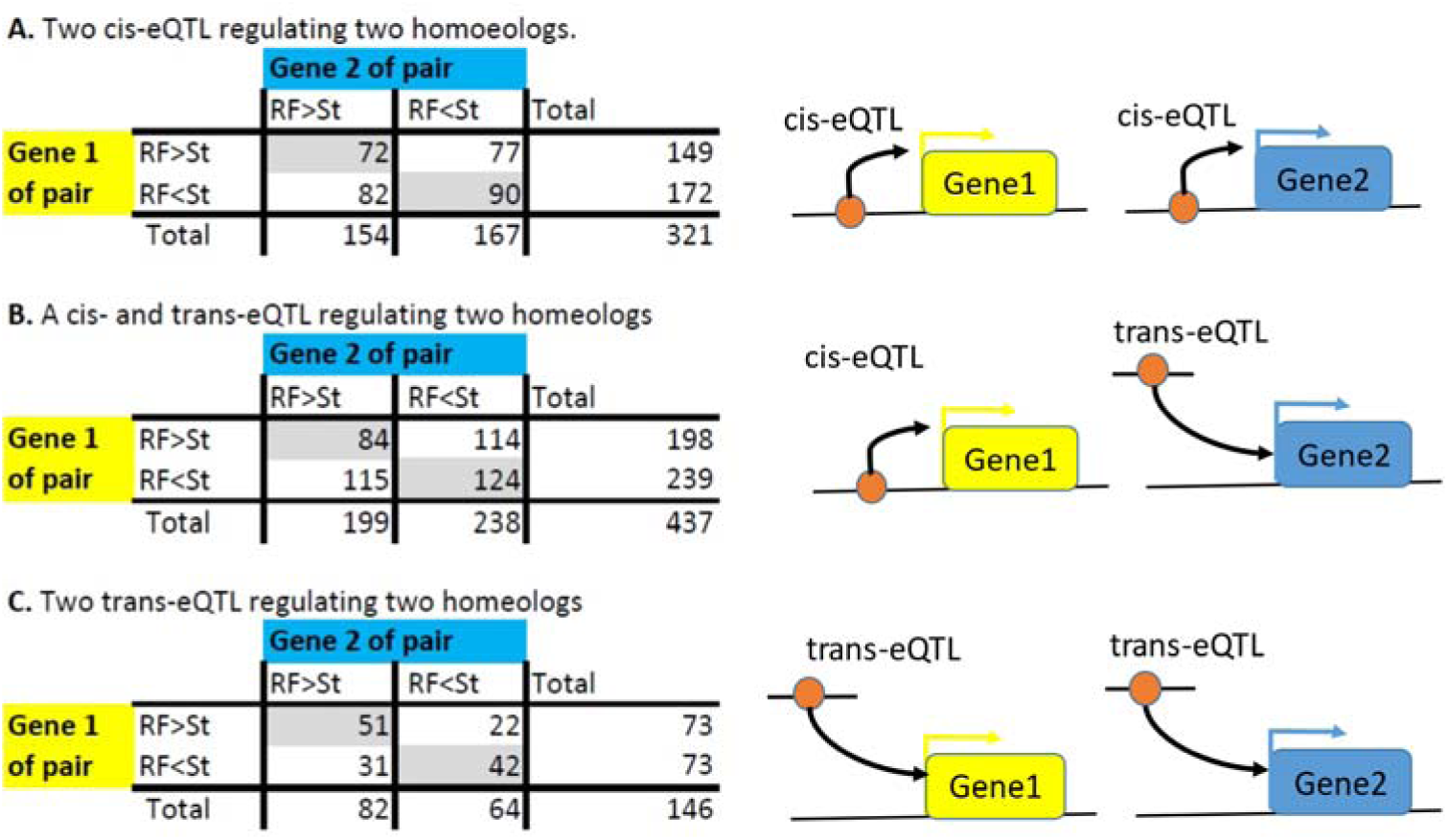
Counts of pairs of homoeologous genes in triplets regulated by two different eQTL and subdivided by parental allelic effects. Gene pairs consistently affected by parental eQTL alleles are given in the grey shaded cells. Gene pairs differentially affected by parents’ eQTL alleles are given in colorless cells. **A** The counts of two genes within a triplet that were regulated by two different cis-eQTL, Type I. **B** The counts of two genes in a triplet whereby one gene was cis-regulated and another gene was trans regulated by separate eQTL, Type II. **C** The counts of gene pairs that were regulated by two different trans-eQTL, Type III. Trans eQTL with the same parental alleles often consistently regulated their target genes (X^2^ test, p-value < 0.001).

Positive selection for different trans-eQTL that co-regulated homoeologous genes suggested that there could have been positive selection for single trans-eQTL that co-regulated homoeologous genes. Co-regulation of duplicate genes was the norm. Eighty-three single trans-eQTL intervals regulated two homoeologous genes, 166 genes total, and nine intervals regulated three homoeologous genes, 27 genes total (**Additional file 1: Table S11; Additional file 2: Fig. S6**).

Alleles consistently co-regulated homeologs in 82 of the 83 target gene pairs, and in all three genes in 3 of the 9 triplets (**Additional file 1: Table S11; Additional file 2: Fig. S6**). While this co-regulation of homeologs was consistent with positive selection favoring single trans-eQTL that had the same effects on homoeologous genes, a level of co-regulation of duplicate genes by a single factor would also have been expected in the absence of positive selection. The transcript levels of different genes bound by the same transcription factor are often highly correlated [44].

The 146 pairs of the Type III trans regulatory loci genotypes notably contributed to grain yields, height, spike length, and thousand kernel weight (Welch Two Sample t-test, P<0.001) (**Additional file 2: Fig. S7**). Variable selection identified six trans eQTL pairs that contributed to yields. Importantly, homozygous Stettler eQTL genotypes at all 12 loci in the six pairs positively contributed to yield (**Fig. 9**). Variable selection also identified five pairs of regulatory alleles that contributed to plant height, and Stettler regulatory genotypes in all five pairs reduced plant height **(Fig. 9)**. Stettler genotypes in 4 of 5 identified pairs increased thousand kernel weight, and Stettler alleles in 3 of 4 identified pairs reduced spike length (**Fig. 9**).

**Fig. 9.**
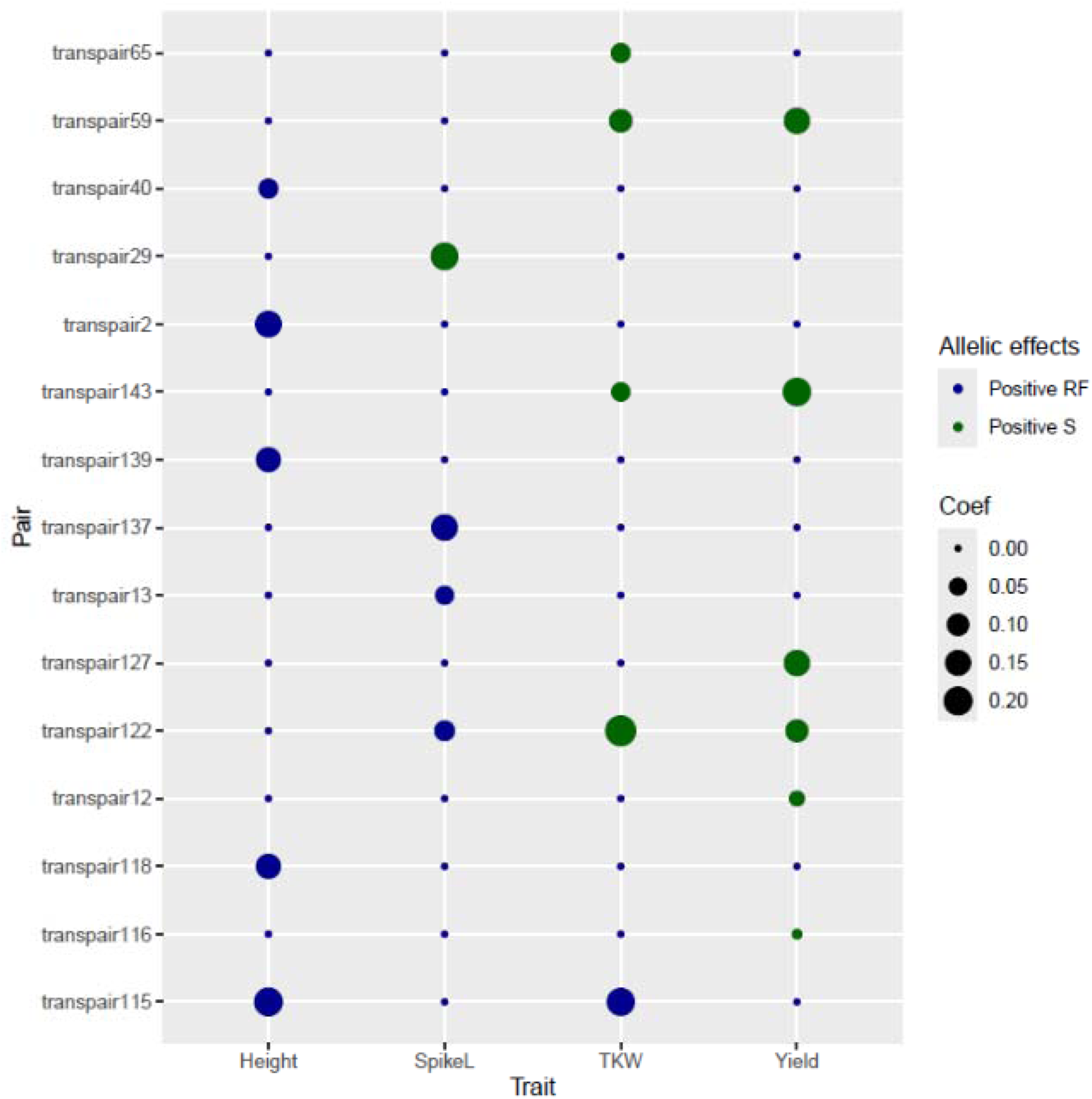
The magnitudes and signs of Type III eQTL pairs on genotypes’ effects on trait values. Effect sizes represent one of two homozygous trans-eQTL’s contribution to the trait. The coefficient estimate, the effect of a single, homozygous trans eQTL, is expressed in standard deviations of z-transformed traits. Trans-eQTL pairs for which homozygous Red Fife trans-eQTL genotypes increased a trait were colored dark blue. Trans-eQTL pairs for which Stettler genotypes increased a trait were colored dark green.

## Discussion

### Many regulatory loci between old and new bread wheat cultivars regulated shared genes and genes on homoeologous chromosomes

Old and modern wheat cultivars had many cis and trans acting regulatory polymorphisms. A total of 12,514 regulatory loci, roughly equally distributed between cis and trans acting loci, affected 10,949 of the 30,296 variable expressed genes (36.1%) (**Fig. 1**). This level of regulatory genetic variation is remarkable because Red Fife is an ancestor of Stettler. Both cultivars are members of the same market class, adapted to the same target environment, and are genetically similar relative to other Western Canadian wheat cultivars [45]. Many more regulatory loci likely differentiate the cultivars. Genes with no identified eQTL usually had higher expression variances across the 154 offspring genotypes than within replicates of the same genotypes. Analysis of a larger population would likely capture additional small effect alleles, likely enriched for trans-acting loci. In an analysis of 112 yeast lines generated from two distinct parents, on average fewer than one eQTL regulated each gene [46]. A later analysis of 1,012 lines from the same population detected close to six eQTL per gene [47].

Different trans-eQTL often regulated the same genes. The number of trans-eQTL was 5,878, and a trans-eQTL containing genetic interval regulated a median of eight genes (**Fig. 2B**). Nonetheless, only 5,328 distinct genes were regulated by a trans-eQTL (**Additional file 1: Table S1**). The number of trans-eQTL similarly outnumbered the number of trans-eQTL regulated genes across diverse bread wheat germplasm [15] and in a maize biparental population [48]. Different stresses often affect the same genes’ transcript abundances [49]. Trans-eQTL likely shared gene targets because different eQTL regulated the same or overlapping abiotic and biotic stress response pathways (**Fig. 6**).

Trans-acting regulatory polymorphisms often affected genes on homoeologous chromosomes. While ten percent of trans eQTL targets were expected to be on homoeologous chromosomes, 25% were (**Fig. 3B; Additional file 1; Table S7**). This enrichment was in part due to one homeolog of a triplet compensating for an expression change at another homeolog, indicating a feedback mechanism to maintain total gene expression levels **(Fig. 4)**. This result reinforces previous findings in polyploid oilseed, *Brassica napus*, and cotton, *Gossypium hirsutum* [15, 16].

Previous studies in wheat and coffee [17, 18] demonstrated that regulatory pathways preferentially targeted genes on specific sub-genomes. While we expected that genes differentially expressed due to Red Fife and Stettler regulatory polymorphisms would be sub-genome enriched, an enrichment was only rarely observed. Trans-eQTL targets taken together were evenly distributed across sub-genomes (**Fig. 3A; Additional file 1: Table S7**), and genes from only three percent (4/119) of the map intervals that regulated 10 or more unlinked genes were enriched in a sub-genome (FDR p<0.05; **Additional file 1: Table S5**). Regulatory polymorphisms may have rarely preferentially affected a sub-genome because our population was not sampled after exposure to abiotic or biotic stress as in the previous reports. Growth in these conditions may have revealed polymorphic regulatory loci with sub-genome specific targets. Second, previously reported sub-genome enrichment may have been due to regulatory loci that preferentially regulated genes on homoeologous chromosomes. When evaluating sub-genome enrichment of trans-eQTL targets, we excluded genes on chromosomes homoeologous to the chromosome containing the trans-eQTL. Including these genes revealed more sub-genome enriched targets (**Additional file 1: Table S4**). Regulatory loci themselves were unequally distributed across sub-genomes. The D sub-genome had a small number of polymorphic regulatory loci compared to the other sub-genomes. Reflecting the pattern in diverse germplasm [50], the Western Canadian breeding germplasm’s D sub-genome was previously reported to be less diverse relative to its A and B genomes [45].

### Selection acted on many regulatory alleles, and its strength depended on target gene functions

We identified high numbers of eQTL relative to synonymous site polymorphisms in low-recombining chromosome central regions relative to high-recombining chromosome arms (**Fig. 5**). Low recombination limits the effectiveness of positive selection to fix beneficial variants and the effectiveness of purifying selection to eliminate deleterious variants [30]. Thus, this result indicated that regulatory polymorphisms had affected fitness. The enrichment of Red Fife and Stettler nonsynonymous polymorphisms in chromosome central regions was similar to the enrichment of eQTL suggesting that regulatory and coding sequence mutations had similar fitness effects. Interestingly, Ganko et al. (2007) reported a strong positive relationship between expression divergence and nonsynonymous substitutions between ancient polyploid-derived gene pairs in *Arabidopsis thaliana* [51] indicating duplicated genes’ regulation and coding sequences may diverge similarly over short and long timescales. Selection in wild populations prior to domestication likely generated the distribution of regulatory polymorphisms between Red Fife and Stettler. Wheat domestication occurred less than 10,000 years ago, and most bread wheat alleles were inherited from wild, ancestral populations [50, 52]. Nonetheless, artificial selection on alleles would also be more efficient at chromosome ends and lead to an enrichment of eQTL in central chromosome regions.

Past selection on regulatory alleles may help explain why trans-eQTL typically had smaller effects on transcript levels than did cis-eQTL. A cis-eQTL explained a median of 46.4% of its single gene target’s transcript variance. A trans-eQTL explained a median of 24.0% of its target gene transcript variance **(Fig. 2A-B)**, but a trans-eQTL regulated a median of 8 genes. Purifying selection has been proposed to preferentially eliminate broadly acting trans-regulatory alleles in natural populations [11, 22, 23]. The trans-eQTL effects also could have been smaller than cis-eQTL effects because the mutations that generated trans-acting regulatory allele had smaller effects on gene expression than the mutations that generated cis-acting regulatory alleles [53].

Genes that were not triplicated had more regulatory polymorphisms between Red Fife and Stettler than did triplicated genes. This result suggested that mutations in non-triplicated genes’ regulatory sites had affected fitness less than mutations in triplicated genes’ regulatory sites. Amongst wheat cultivars, triplicated genes have also been reported to have fewer nonsynonymous polymorphisms than non-triplicated genes [20, 21].

Genes contributing to environmental responses were more likely to have cis-eQTL and especially trans-eQTL than genes contributing to other biological processes **(Fig. 6)**, consistent with selection acting differently on regulatory alleles that contributed to environmental responses compared to other processes. Regulatory alleles affecting environmental responses may have had less of an impact on fitness in both wild and domesticated germplasm than regulatory alleles affecting other processes and thus were maintained in wheat germplasm. More likely, positive selection for environmental response alleles contributed to the high number of regulatory allele differences. Especially prior to modern agronomic practices, past genotypes were subjected to diverse abiotic and biotic challenges, and duplicated genomes have a high capacity to generate adaptive, environmental responses. Polyploid yeast strains grown in a poor carbon source media acquired mutations with high fitness effects and quickly adapted to environmental stress [54], and neofunctionalization of genes following Brassica polyploidization probably allowed plants to escape herbivory [55, 56]. Artificial selection rapidly changed crops’ environmental responses. In the era separating Red Fife and Stettler, breeders have selected for genes increasing grain yield potential and grain yield stability under diverse agricultural environments [29] and for disease resistance genes such as Lr2a [32,57]. Liu et al. (2023) suggested selections under conditions of sufficient fertilizer supply caused current *Brassica napus* oilseed varieties to have less developed roots, fewer root exudates, and lower yields when grown under low phosphate conditions [58]. Ma et al. (2019) reported the number of disease resistance genes in cultivated barley was low compared to wild barley and suggested the difference was due to the physiological cost of maintaining the alleles [59].

### Plant breeders selected for hotspot regulatory alleles

Artificial selection for hotspot regulatory alleles contributed to wheat improvement. The Stettler genome had two alleles that were known selection targets, the *Rht-B1b* semi-dwarfing allele and the *Lr2a* rust resistance allele. Both alleles generated regulatory hotspots **(Fig. 7)**. In addition, allelic variation at the 42 hotspot regulatory loci significantly contributed to yield and other traits targeted by plant breeders. Stettler hotspot alleles were much more likely to contribute favorably to traits than were Red Fife alleles. Among the six yield-contributing hotpots, Stettler genotypes at five increased yields (**Fig. 7)**. Hotspot regulation of diverse pathways contributed to greater yield potential and yield protection **(Additional file 1: Table S10)**. Transcription factor differences likely accounted for some hotspots. Transcription factors associated with hotspots in yeast [47], and transcription factor changes have generated domestication and improvement phenotypes in many plant species [6, 60]. Nonetheless, differences in other genes can affect physiological and developmental processes that in turn generate extensive expression changes [47]. For example, hotspot Rht-B1 encodes a DELLA protein that functions in the GA signaling pathway.

We expect polyploidization and cultivation contributed to the favorability of hotspot alleles. For example, *Rht-B1b* is beneficial in an agricultural system but would be deleterious in natural environments. Other favorable hotspot alleles such as the five Stettler yield-increasing loci (**Fig. 7**) may also have positive effects only in cultivated environments. We would expect these alleles to be absent or at low frequencies in ancestral wheat germplasm [61]. Hotspots may be also favorable because other homeologs buffered their negative effects by maintaining their ancestral functions [19, 24]. Evaluating lines with Stettler hotspot alleles with and without mutagenized homeologs would test this idea.

### Plant breeders selected different trans eQTL alleles that consistently regulated homeologs

The accumulation of trans-eQTL with consistent effects on homoeologous genes in the modern cultivar Stettler suggested plant breeders selected for similar expression patterns in homoeologous genes in the same triplet. When two trans-eQTL regulated different homeologs, Type III pairs, reinforcing trans-acting alleles were detected notably more often than expected (**Fig. 8**). These regulatory pairs contributed to yield and other traits significantly more than expected by chance **(Additional file 2: Fig. S7)**. Variable selection identified six homeolog pairs, 12 loci, that contributed to yield, and lines with Stettler alleles at all six pairs had superior traits compared to lines with Red Fife alleles (**Fig. 9**). The coordinated regulation of homeologs by separate regulatory loci is noteworthy. The single regulatory alleles that affected two or more homeologs within a triplet also had consistent effects (**Additional file 1: Table S11; Additional file 2: Fig. S6**).

Plant breeders generated new varieties by selecting inbred or doubled haploid offspring from biparental crosses. If several favorable loci differed between parents in a cross, very few offspring genotypes would contain a large number of the favorable regulatory alleles. For yield, the genotypic effects of each of the 12 regulatory loci in the six homoeologous pairs were additive (**Fig. 9**). Thus, favorable alleles regulating both genes in a homoeologous pair likely accumulated in genotypes one or two at a time as wheat germplasm changed over the years [45]. Seed of parental lines used in past breeding crosses are available, and genotyping these lines could determine if beneficial regulatory alleles accumulated rapidly in response to artificial selection. Interestingly, genes retained in polyploid genomes over long time scales were maintained because each copy increased plant fitness [62], again suggesting mechanistic similarities with the evolution of natural populations over the long term and rapid evolution during plant breeding.

We found no evidence that selection favored homeolog reinforcing alleles at two cis regulatory loci, Type I regulation **(Fig. 8A)**, nor at one cis and one trans regulatory locus, Type II regulation **(Fig. 8B)**. One reason may have been that a single cis allele usually had strong effects on target gene expression **(Fig. 2)**, and reinforcing its effects using homoeologous loci may not have been beneficial. He et al. 2022 report a negative correlation between marker allele frequency and cis-eQTL effect size within a diverse wheat population [15]. Pyramiding cis-regulatory alleles at different homeologs also appears to rarely generate favorable traits. In a diverse set of wheat genotypes, while there was evidence of selective sweeps on different homeologs, one individual rarely had more than one homeolog with a selected allele [20]. Similarly, You et al. (2023) explored gene expression variation in developing tetraploid cotton fibers across a genetically diverse population [63]. When one homeolog favorably contributed to a fiber trait, the second homeolog rarely contributed. QTL effects on a target trait can be less-than additive. In sugarcane, genetic variation at duplicated genomic regions contributed to sugar content variation. However, a single region generated most of the desired effect on sugar content. Adding additional regions generated small responses [64].

## Conclusion

This work identified thousands of regulatory locus polymorphisms that generated gene transcript abundance differences between old and new hexaploid bread wheat cultivars. Allelic differences at trans-eQTL preferentially affected genes on homoeologous chromosomes but largely equally affected genes across sub-genomes. Continued crop improvement is an important goal. Results indicated that despite the presence of homoeologous genes, selection on regulatory alleles was widespread. Thus, the large number of genes in polyploid wheat provide raw material for mutation to generate favorable, novel regulatory traits, especially for environmental response genes. Results also suggested that new mutations may be favorable because of genome duplication. Breeders selected both hotspot regulatory alleles which can be buffered by homeologs and different trans-acting regulatory alleles that targeted duplicated genes. The capacity to generate favorable genetic variation may help explain the past success of polyploids and point to future breeding gains.

## Materials and Methods

### Germplasm, plant growth conditions, and tissue harvesting

To understand the frequency and attributes of wheat gene regulatory variants and to investigate selection on gene regulation, we identified the regulatory allele differences between an old and a new wheat cultivar. Gene transcript abundances were estimated across a doubled haploid (DH) population of 154 lines derived from the cross between Red Fife and Stettler hard-red spring wheat cultivars. Red Fife was developed in 1841, has excellent bread-making quality [57], and set Canadian wheat standards. It was the dominant cultivar in Western Canada until the early 1900s [57] and is still grown today for organic and specialized markets. Red Fife is an ancestor of Stettler that was released in 2008 [65], and Stettler has high grain yield, protein content, lodging resistance, and stem rust resistance [65]. It is more disease resistant than Red Fife, and its yield is much higher [21]. Stettler was grown on 106,105 hectares in 2020 in Western Canada (https://grainscanada.gc.ca/en/grain-research/statistics/varieties-by-acreage/). Genetic analysis of a homozygous population derived from a biparental cross has a number of advantages relative to the analysis of diverse germplasm. Linkage disequilibrium between a marker and a trait unambiguously indicates linkage; lines within the population have not been subjected to intentional selection; there is high power to detect allelic effects because the expected frequency of each polymorphic allele is 1:1; and larger linkage blocks reduce the number of statistical tests.

For RNA extraction, Red Fife, Stettler and their DH lines were grown in a growth room as described in Raherison et al. 2020 with three replicates in time [21]. For each replicate, Red Fife and Stettler were grown in four pots, and each of the 154 DH lines was grown in a single pot. The second leaf from the plant base was harvested at the same day and time across 12-day-old growth room plants (Zadoks stage Z1.2) to ensure developmental and circadian stage uniformity across lines. For each line, tissues across the three replicates were combined into one sample for RNA extraction. Thus, there was one pooled tissue sample per DH population line and four pooled tissue samples per parental line.

### RNA extraction, RNA-seq, and eQTL mapping

Total RNA was extracted from each sample using an in-house Trizol-based method. Poly-A containing mRNA molecules were purified, and libraries of 200-bp insert sizes were constructed according to the Illumina TruSeq RNA sample preparation guide v2. RNA-seq was performed on each sample using the HiSeq 2500 platform at the University Health Network Clinical Genomics lab in Toronto, Ontario.

FastQC 0.11.9 [66] assessed read quality, and Trim Galore 0.6.6 [67] trimmed reads and removed low-quality nucleotides. Trimmed reads shorter than 75 bp were removed. On average, each sample had 20.06 million processed 100-125 bp long paired end reads comprising a total of 3.25 billion high-quality paired end reads.

The kallisto/0.46.1 software assigned the reads to transcripts from the Chinese Spring RefSeqv1.0+UTR annotated transcriptome. The kallisto/0.46.1 software had accurate homoeolog-specific read mapping in hexaploid bread wheat [68]. The tximport/1.18.0 R package converted kallisto’s transcript level counts to gene level counts [69] across the 154 DH samples and the 4 parental replicates.

CoGe analysis (https://genomevolution.org) classified the 110,790 high-confidence genes in the annotation into one of three categories [21]. A total of 43,254 genes were categorized as members of a unique triplet. Each gene of a unique triplet matched to a single gene on each of the two remaining homoeologous chromosomes at homoeologous locations. Unique triplet groups numbered 14,418. A total of 10,945 genes were categorized as members of 11,814 non-unique triplet groups. A non-unique triplet gene had homoeologous genes in the two other sub-genomes, but duplications within a sub-genome caused some genes in a triplet to also be classified in another triplet. Our analysis focused on unique triplet genes. The remaining 56,591 genes, non-triplicated genes, were classified in the non-triplet group. Each gene in this group did not match to a gene on each of the two remaining homoeologous chromosomes at homoeologous locations.

Of the 110,790 annotated genes, we selected 40,395 genes that had > 1 count per million paired reads in at least 65 samples, 40% of all samples. EdgeR-3.32.0 normalized read counts using the trimmed mean of M-values (TMM) approach and log transformed gene expression values [70]. Principal components (PCs) adjusted the log-transformed TMM data. When only parental lines’ TMM data was analyzed, the four samples from each parent had similar PC1 values. In a PC analysis of all lines’ TMM values, some parental samples grouped closer to DH lines than to each other parental samples. We regressed each gene’s TMM values from all samples on the first three PCs to correct for noise such as library and/or sequencing batch effects and used the residuals as the estimated gene expression values [71]. Parental samples clustered in the PC analysis of this corrected data. The top 75% most variable genes (30,296) across the 154 DH lines were used for eQTL mapping.

We mapped eQTL to genetic map positions so as to investigate eQTL inheritance and attributes related to crossover frequencies. A genetic map with 1,483 SNP sites across 25 linkage groups corresponding to the 21 wheat chromosomes was previously reported [21]. R/qtl composite interval mapping with default settings mapped eQTL [72]. 1000 permutations were generated to estimate genome-wide LOD thresholds at a 5% significance level. An eQTL that was on the chromosome of its regulated gene was classified as a cis-eQTL. An eQTL that was on a different chromosome from its regulated gene was denoted as a trans-eQTL. We used 8.8cM as the 95% confidence interval for a trans-eQTL’s location because a trans-eQTL LOD score was reduced by an average of 1.5 LOD 4.4cM away from its LOD peak [73].

### Identifying genes regulated by the same eQTL interval and their sub-genome locations

We calculated the number of genes affected by trans-acting regulatory loci using two methods. First, we identified trans-eQTL hotspots, single chromosomal intervals that regulate many genes, by counting the number of eQTL peaks in 3cM genetic map intervals using a 1cM step-distance. Hotspots were identified if a single 3cM interval contained at least 20 trans-eQTL peaks. Hotspots were extended if a neighboring 3cM interval also contained at least 20 trans-eQTL peaks. Second, we counted the number of genes that had trans-eQTL peaks within each 8.8cM interval on the genetic map. As noted in the text, genes affected by different, closely linked regulatory sites would have trans-eQTL in the same hotspot or interval.

To investigate if trans-eQTL target genes were uniformly distributed across the A, B and D sub-genomes, we recorded the locations of trans-eQTL target genes. Sub-genome specificity was tested with the X^2^ test. We also identified the genetic map intervals within which at least 10 genes had trans-eQTL and evaluated the probability that these co-regulated genes were randomly distributed across sub-genomes using the X^2^ test. The Benjamini-Hochberg procedure controlled the FDR to 0.05. We excluded target genes on homoeologous chromosomes from this sub-genome analysis since an eQTL on one homeolog preferentially regulated genes on other homeologs, as described in results. We tested if genes on homeologs were enriched using one-sample proportions tests.

### Tests for regulatory allele selection in ancestral and plant breeding populations

To infer if selection had acted on eQTL, we determined if eQTL accumulated in central, low-recombination chromosomal regions, where selection is relatively ineffective relative to terminal, high recombination regions [30]. The eQTL counts in each region were adjusted by the number of synonymous SNPs identified by SnpEff v4.3T [21], and the number of expressed genes in the region. Bedtools/2.29.2 tallied both. Central regions with low crossing-over frequencies were delimited for each chromosome by eye using plots of SNPs’ genetic map positions relative to their physical map positions. Differences in eQTL number between chromosomal regions were compared using paired t tests. To compare the frequency of eQTL with the frequency of nonsynonymous differences, we evaluated the ratio of non-synonymous to synonymous SNPs across regions corrected for the number of expressed genes, also using paired t-tests.

We tested if gene ontology terms were unequally enriched in sets of eQTL regulated genes. A set of 28,081genes of the 30,296 variably expressed genes were annotated with GO terms. We compared each set of eQTL regulated genes of interest with the 30,296 most variable genes as the gene universe for GO term enrichment using topGO/2.48.0 [74] with an alpha value of 0.01.

Plant breeders’ selections may have favored consistent up- or down-regulation of genes within a triplet. Analysis of genes within the same triplet regulated by separate eQTL can test this idea. An abundance of parental alleles at different eQTL that have similar effects on their target genes indicate selection. We examined pairs of triplicated genes in which each gene of a pair was regulated by different eQTL. Three types of triplet gene pairs were analyzed. In Type I, two cis-eQTL regulated the two genes. In Type II, one trans-eQTL and one cis-eQTL regulated the two genes. In Type III, two trans-eQTL regulated the two genes. For each group, Chi Square tests evaluated if parental eQTL effects on genes within a triplet were independent.

To determine if predicted selection targets affected important traits, eight agronomic traits for the DH lines and their parents were measured on field-grown plants. The experimental design was a randomized complete block design at two locations with two blocks at each location. Line values were estimated using a linear model. Cultivation and trait details were described in Raherison et al. 2020 [21]. The eight agronomic traits were days after planting when 50% of plants within a plot had headed (Head50); days after planting when 100% of plants within a plot had headed (Head100); the distance from the ground to the end of the spike after heading (Height); plant lodging (Lodging); protein content of the grain (Protein); spike length (SpikeL); thousand kernel weight (TKW); and yield (Yield) [21].

LASSO (Least Absolute Shrinkage and Selection Operator) regression determined if the Type III eQTL pairs contributed to traits. Trait data was z transformed, and individual lines’ trait values were modeled as a linear function of the number of homozygous Stettler genotypes within each of the eQTL pairs. Each eQTL pair was considered a single factor with three levels: −1, if the line was homozygous for Red Fife alleles at the two trans-eQTL, 0 if the line was homozygous for one Red Fife allele and one Stettler allele at the two trans-eQTL; and 1 if the line was homozygous for two Stettler alleles at the two loci. The model was fitted using glmnet 4.1-8 [75]. To evaluate if observed LASSO results could be generated in the absence of a phenotype-genotype association, mean square cross-validation errors of 100 LASSO regression models repeatedly fit on the data were compared with mean square errors of 100 LASSO models in which trait values were randomly associated with lines. Those pairs of homoeologous genes’ trans-eQTL selected by LASSO regression were retained. Coefficient estimates, representing the effect of a homozygous Stettler genotype at one eQTL locus, were reported in terms of standard deviations.

Breeders’ selection on hotspot intervals was evaluated by cross-referencing hotspot intervals with known genes selected by wheat breeders and by using LASSO regression to determine if hotspot alleles notably contributed to plant improvement traits. We did a literature review to identify favorable genes that were unambiguously present in Stettler and absent from Red Fife. The regression of yield and other traits on hotspot genotypes was done in a similar way as described for eQTL pairs except the hotspot loci were predictors. Each hotspot locus was coded as a factor with two levels −1 for homozygous Red Fife and 1 for homozygous Stettler. The coefficient estimates represented one-half of the difference between homozygote trait values.

## Abbreviations

RNA-seq: RNA sequencing;
SNP: Single Nucleotide Polymorphism
cis-eQTL: cis-acting expression quantitative trait locus
trans-eQTL: trans-acting expression quantitative trait locus
UTR: Untranslated region
DHLs: Doubled Haploid Lines
TMM: trimmed mean of M-values
FDR: false discovery rate
GO: Gene Ontology
BP: biological process
SpikeL: spike length
TKW: thousand kernel weight
GA: gibberellin
LASSO: Least Absolute Shrinkage and Selection Operator.

## Declarations

### Ethics approval and consent to participate

Not applicable.

### Consent for publication

Not applicable

### Availability of data and materials

Raw RNA-seq data in this study can be obtained through BioProject ID [#] on the NCBI Sequence Read Archive [76]. The scripts for analyzing raw data sequences and intermediate data are available under MIT license at GitHub:[77] and in Zenodo or figshare: [78].

These references [76–78] will be given in the reference list if the manuscript is accepted. Dataset identifiers including DOIs will be expressed as full URLs.

### Competing interests

The authors declare that they have no competing interests.

### Funding

This research was funded by Genome Canada and the Natural Sciences and Engineering Research Council of Canada.

### Authors’ contributions

RC, RK: Generated population.

ER, MH, NH: Grew plants, measured traits, isolated RNA.

ER, MH, MM, RG, SZ, WH: Analyzed data.

SZ, LL: Wrote manuscript.

## Acknowledgements

We thank our colleagues for our discussions about this work. We also thank the Digital Research Alliance for providing computing infrastructure.

## Notes

### Competing Interest Statement

The authors have declared no competing interest.

