## Additional File 2: Supplementary Figures S1-7 for "Positive selection on hotspot and reinforcing regulatory alleles contributed to hexaploid bread wheat improvement"

**Additional File 2: Supplementary Figures for “Positive selection on hotspot and reinforcing regulatory alleles contributed to hexaploid bread wheat improvement.”**

Shuhua Zhan,^1^ Elie Raherison,^1^ William Hargreaves,^1^ Nia Hughes,^1^ Roos Goessen,^1^ Mohammad Mahdi Majidi,^2^ Ron Knox,^3^ Richard Cuthbert,^3^ and Lewis Lukens^1^*

^1^Department of Plant Agriculture, University of Guelph, Guelph, Ontario, Canada

^2^Department of Agronomy and Plant Breeding, College of Agriculture, Isfahan University of Technology, Isfahan, 84156-83111, Iran

^3^Swift Current Research and Development Centre, Agriculture and Agrifood Canada, Swift Current, Saskatchewan

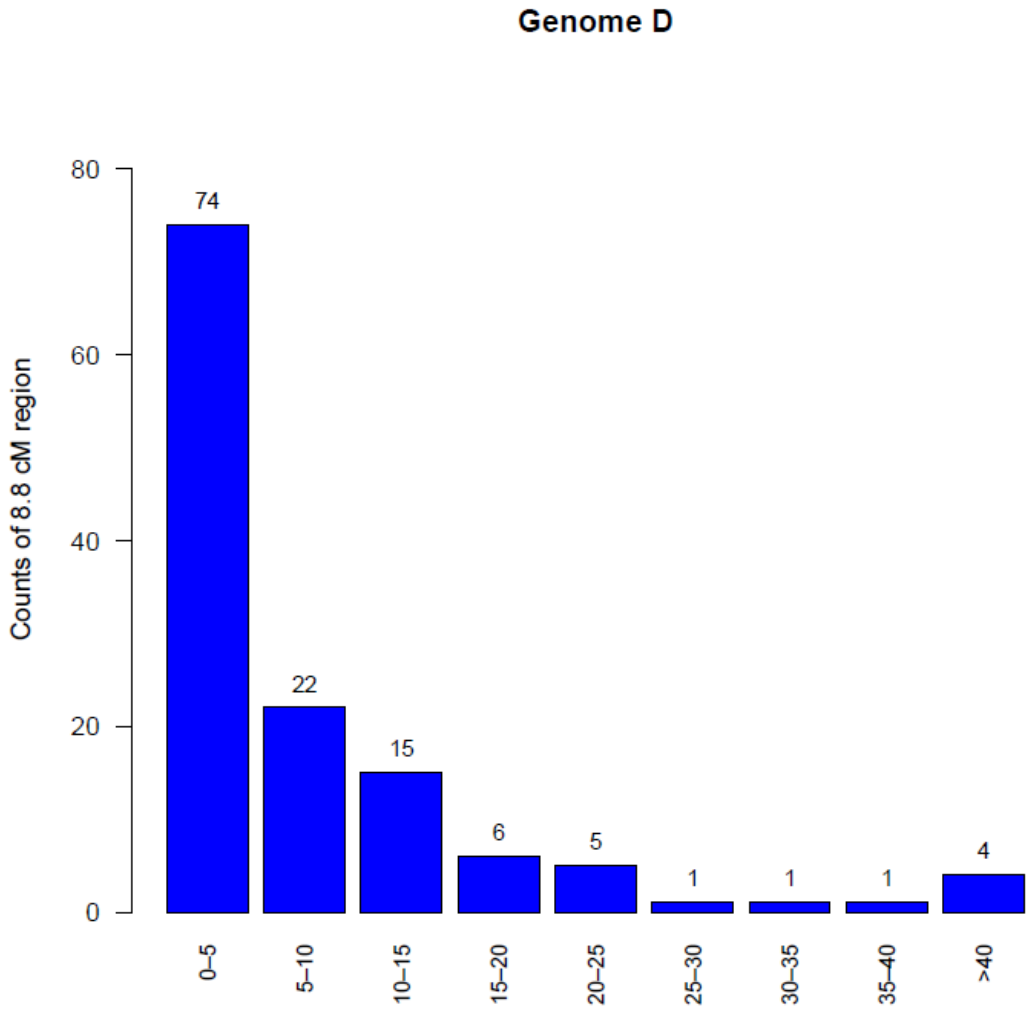

**Fig. S1** The number of genes with eQTL that map to the same 8.8 cM interval on the D genome.

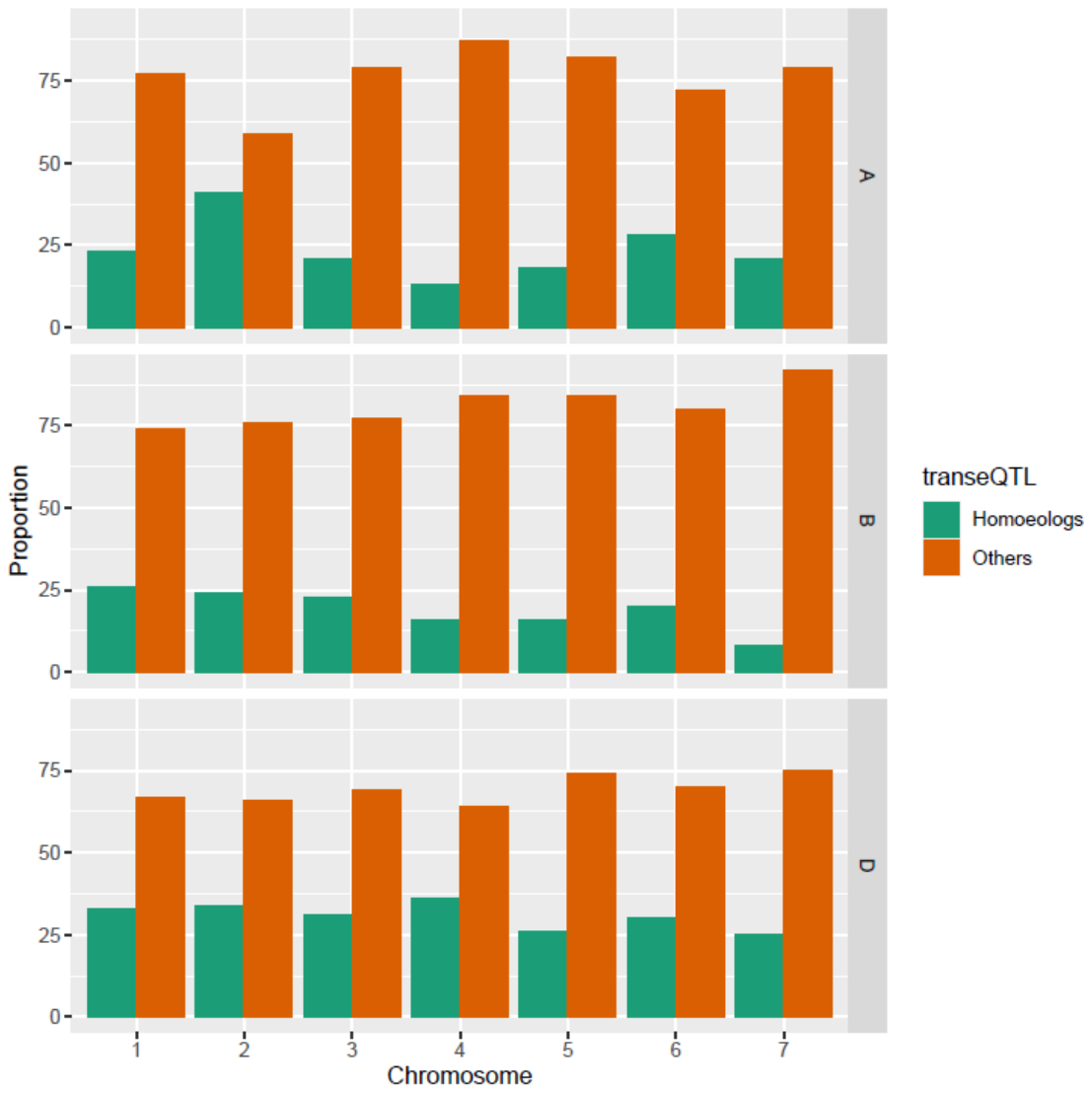

**Fig. S2** Proportions of each chromosome’s genes regulated by trans-eQTL on homoeologous chromosomes or other, non-homoeologous chromosomes.

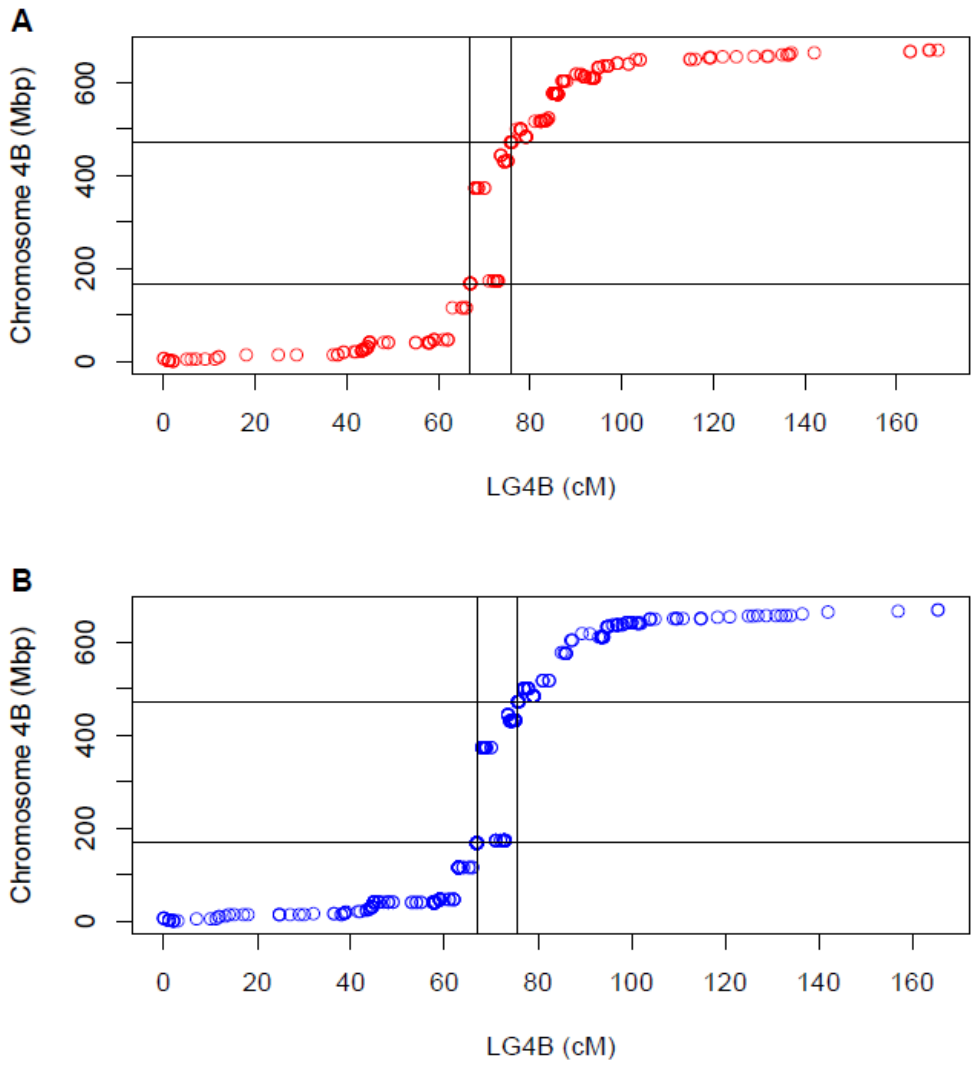

**Fig. S3** Recombination was much lower in the center of chromosomes relative to chromosome ends. A scatter plot of 4B cis-eQTL (**A**) and trans-eQTL (**B**) genetic and genomic positions. Horizontal and vertical lines demarcate the large central region with notably low recombination.

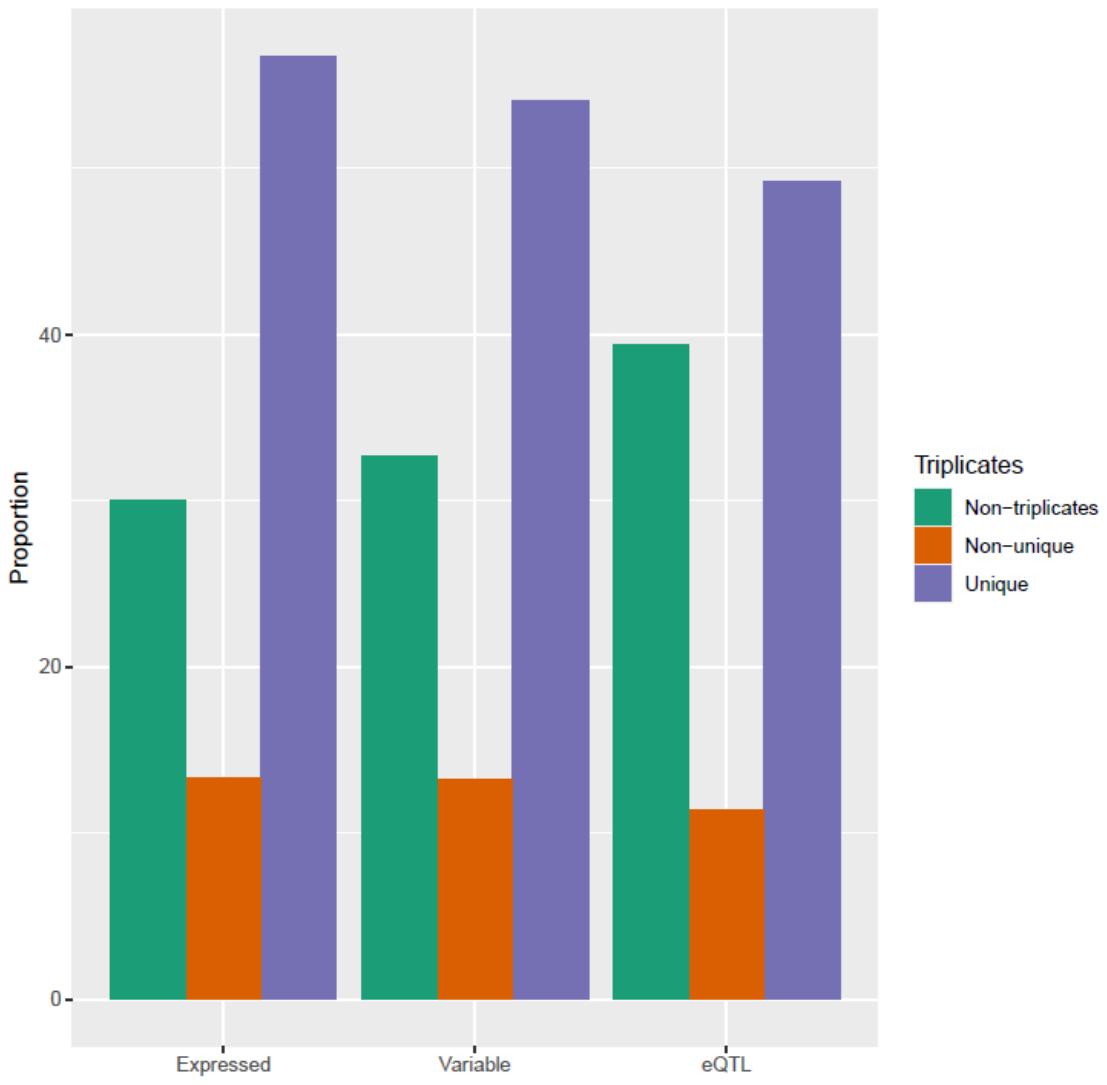

**Fig. S4** Triplicated genes’ variability and eQTL regulation is lower than non-triplicated genes. The proportion of variable genes that are triplets is less than the proportion of expressed genes, and the proportion of eQTL regulated genes is lower than the proportion of regulated genes.

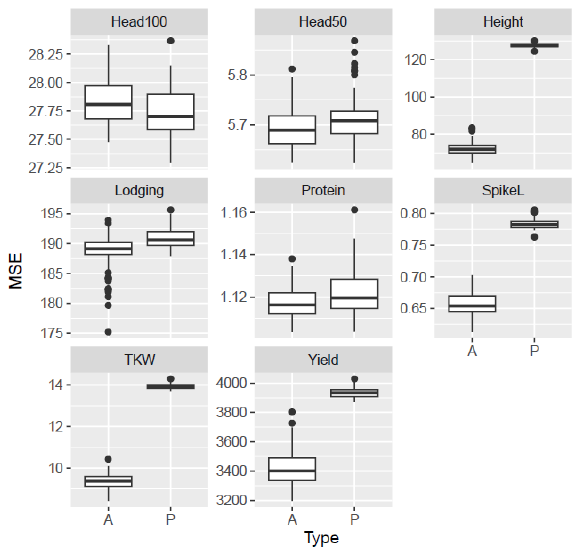

**Fig. S5** Comparison of regression models fit on actual (A) and permuted (P) hotspot eQTL genotype and trait datasets. Mean squared errors (MSEs) of hotspot predicted traits were plotted from 100 LASSO analyses of the eight traits using 42 hotspot eQTL as predictors across actual and simulated data sets. For height, spike length, TKW, and yield, actual datasets generated notably lower MSEs than the permuted datasets. Not one model fit on permuted trait values generated a cross-validation error lower than a fit on actual data.

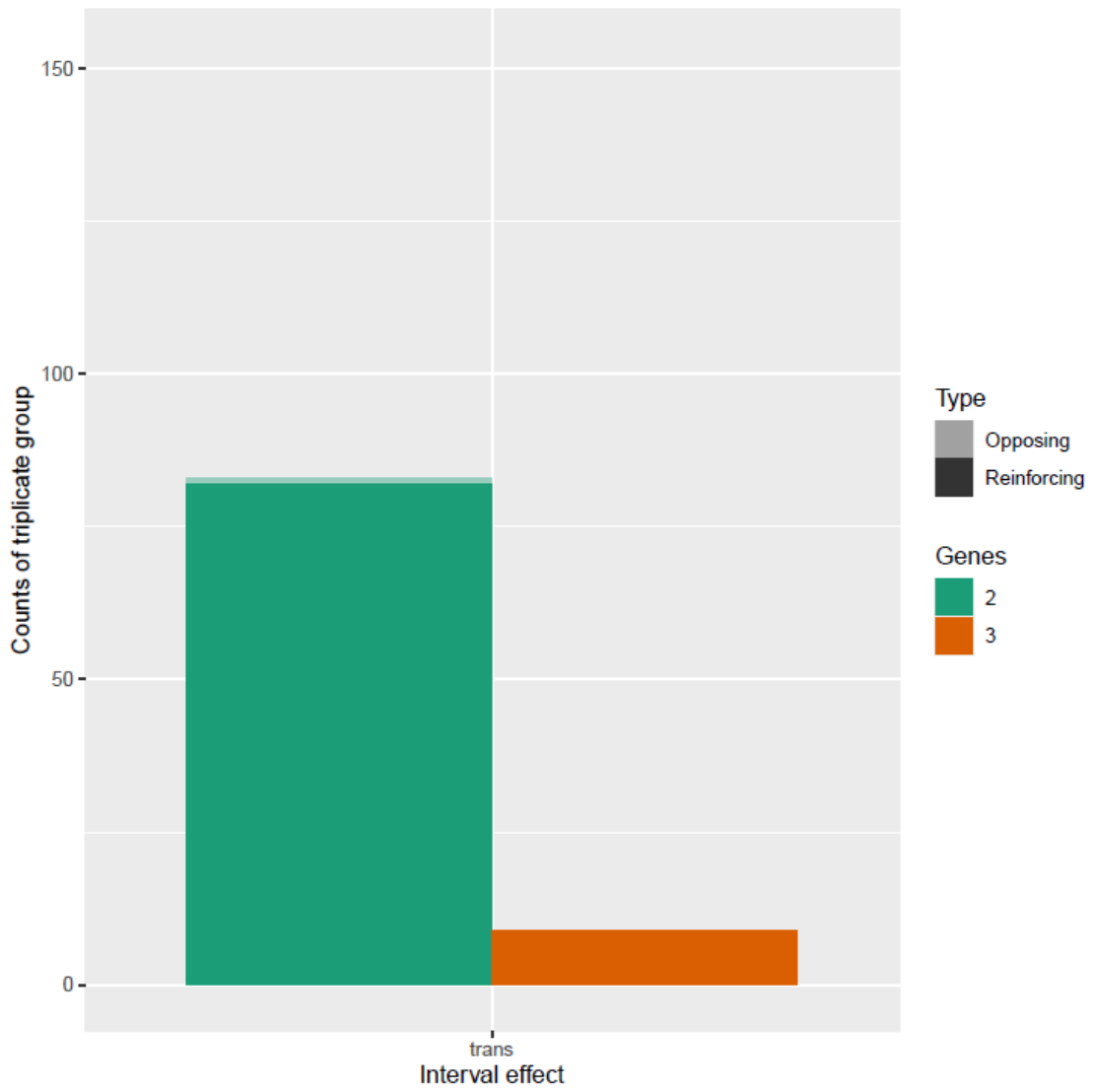

**Fig. S6** eQTL regulation and co-regulation of triplicate genes. The number of gene pairs within triplicate groups in which all homeologs were trans regulated by the same locus is plotted. A single Stettler trans-eQTL had consistent effects on close to all duplicated genes.

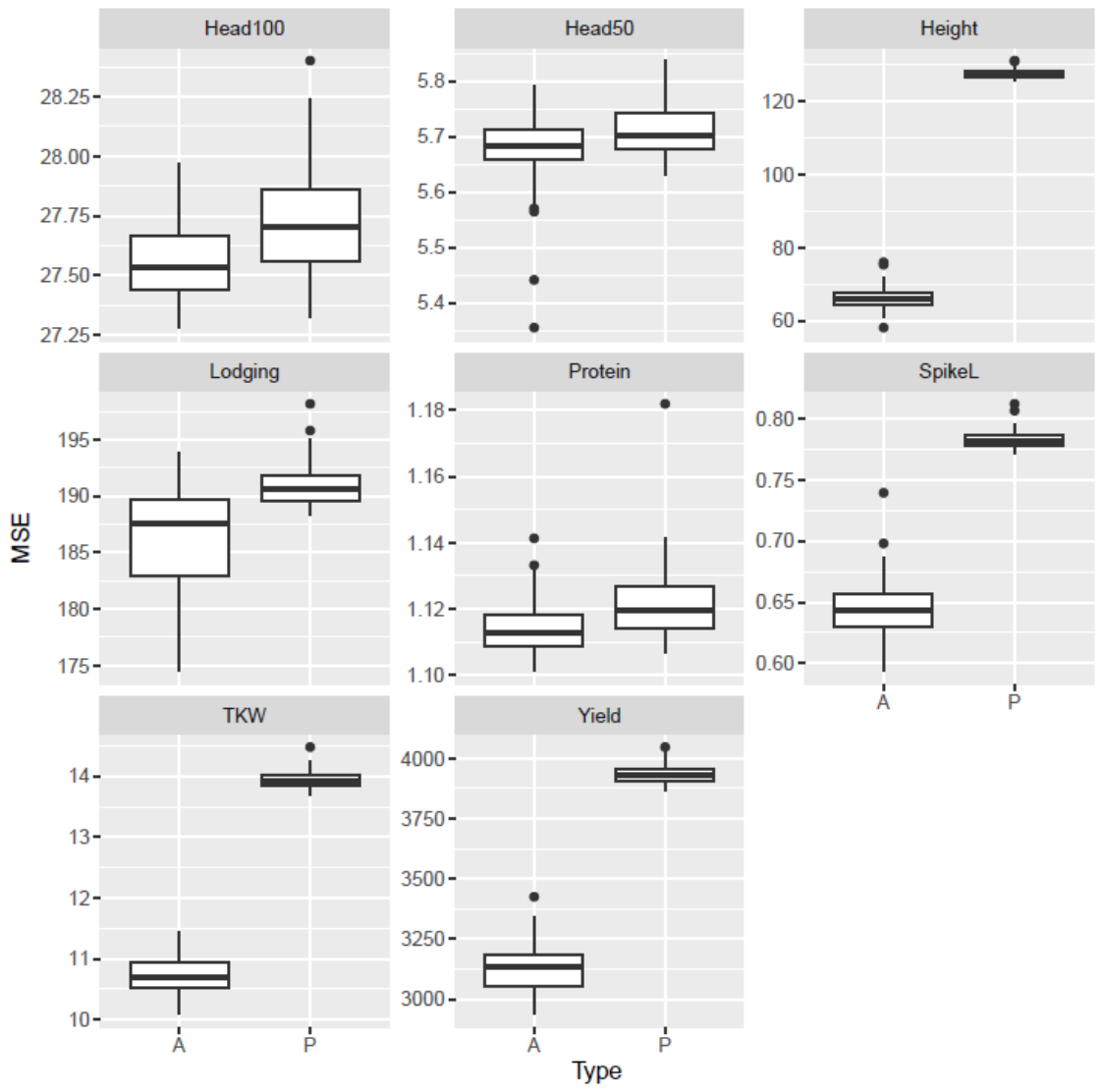

**Fig. S7** Comparison of regression models fit on permuted (P) and actual (A) datasets that paired homoeologous genes’ trans-eQTL genotypes and traits. Mean squared errors (MSEs) were plotted from 100 LASSO analyses of eight traits. For height, spike length, TKW, and yield, models fit with the actual datasets generated significantly lower MSEs than those fit with permuted datasets.
